# Human spinal cord organoids exhibiting neural tube morphogenesis for a quantifiable drug screening system of neural tube defects

**DOI:** 10.1101/2020.12.02.409177

**Authors:** Ju-Hyun Lee, Hyogeun Shin, Mohammed R. Shaker, Hyun Jung Kim, June Hoan Kim, Namwon Lee, Minjin Kang, Subin Cho, Tae Hwan Kwak, Jong Woon Kim, Mi-Ryong Song, Seung-Hae Kwon, Dong Wook Han, Sanghyuk Lee, Se-Young Choi, Im Joo Rhyu, Hyun Kim, Dongho Geum, Il-Joo Cho, Woong Sun

## Abstract

The human spinal cord forms well-organized neural circuits for environment sensing and motor behavior. The three-dimensional (3D) induction of the spinal cord-like tissue from human pluripotent stem cells has been reported, but they often do not mimic morphological features of neurulation and their maturity is limited. Here, we report an advanced 3D culture system for the production of human spinal cord-like organoids (hSCOs) suitable for the scale-up and quantitative studies. The hSCOs exhibited many aspects of spinal cord development, including neurulation-like tube-forming morphogenesis, differentiation of the major spinal cord neurons and glial cells, and mature synaptic functional activities. We further demonstrated that hSCOs platform allowed quantitative and systematic high-throughput examination of the potential risk of neural tube defects induced by antiepileptic drugs. Thus, hSCOs can be used for understanding human spinal cord development, disease modeling, and toxicology screening.

## Introduction

The spinal cord plays critical roles in the neurotransmission of sensory inputs and motor outputs between the brain and the body, the coordination of central pattern generation, and many sensory-motor reflexes. During embryonic development, the spinal cord is formed via neurulation, an early morphogenetic process. Typically, neurulation is mediated by sequential processes including polarized neuroepithelial (NE) cell induction, sheet-like neural plate formation, and folding-based tube morphogenesis^1^. The posterior part of the neural tube develops into the spinal cord containing more than 20 classes of neurons that connect other tissues in the body and establish neuronal circuits governing somatosensation or locomotion^2, 3^. Thus, many human diseases associated with the spinal cord lead to abnormalities in sensorymotor reflexes and autonomic nervous system. Deficiencies in the early neurulation process often lead to neural tube defects (NTDs). As one of the major congenital malformations, NTDs can be caused by genetic, nutritional, or environmental factors. More than 200 genes are known to cause NTDs in mouse models^4, 5^. Human genetic studies associated with NTDs have demonstrated limited correlations with mouse mutations. Most of the information about human NTDs is obtained from retrospective clinical research. The primary risk factors for NTDs are folate deficiency, maternal diabetes, and side effects of antiepileptic drugs (AEDs) during pregnancy^6–9^. A huge gap exists between mouse and human studies. The mechanism by which such factors cause or alter NTD pathology remains primarily unknown.

The access to human embryo/fetus is highly limited owing to the ethical and technical limitations. Thus, *in vitro* replication of important features of human embryonic development via three-dimensional (3D) culture of the organoids derived from human pluripotent stem cells (hPSCs) can lead to new opportunities for investigating human development, including three germ layers patterning, early axial organization, and organogenesis^10–13^. The central nervous system (CNS) organoids are considered valuable model systems to explore the most complex and highly organized human nervous system and neurological disorders^14–19^. A 3D organoid system representing the posterior part of the CNS has been reported^20–24^. Such protocols demonstrate the efficiency of spinal cell type induction, dorsoventral specification, and 3D trunk neuromuscular connections. Although advent of organoids offer a new paradigm in biomedical research and neurodevelopmental biology, batch variations, intra- or inter-organoid variations have limited their use in robust quantification-based drug screening or toxicology tests^17, 25, 26^. Morphological and physiological evaluations of the spinal cord organoid system are in the early stages. Thus, the use of these organoids as a drug screening system requires improvement for reducing the inter- or intra-experimental variations and developing accurate quantification systems.

Here, we report a novel method for producing spinal cord organoids recapitulating neurulation-like morphogenesis. Most of the previous organoid models exhibited neural-follicle or cyst expansion similar to 2D neural rosette formation, which is different from neural tube formation *in vivo,* attributing limitations to the current culture system for NTD disease modeling. Our 3D culture system can be used for the rapid production of a large number of spinal cord organoids, allowing for the quantification of the organoid morphogenesis. The robustness of the method was evaluated with a screening of the AEDs that can cause NTDs, which can offer insights to understand the mechanism of neurulation and the toxicology test for the human NTDs.

## Results

### Protocol for the formation of the human spinal cord organoids (hSCOs)

We established a 3D culture system that recapitulates early spinal cord induction with the morphological events of neurulation (Fig. 1a). Our protocol included three consecutive steps. In the first step, the WNT activator CHIR99021 (CHIR) and the inhibitor of TGF-ß signaling SB431542 (SB) were added to the hPSCs monolayer culture for 3 days to induce neural stem cells (NSCs)^27^. During the 2D induction process, hPSCs were determined as caudal neural stem cells (cNSCs) at the neuromesodermal progenitors (NMps) stage, as observed in embryonic caudal neurogenesis *in vivo*^28^ (Supplementary Fig. 1a-d). Next, cNSC colonies were gently detached from the dish to form 3D sphere. These spheres were cultured in the presence of bFGF for 4 days for the expansion and establishment of the neuroepithelial (NE) alignment (Supplementary Fig. 1e, f and Supplementary Video 1). Lastly, the media was changed to favor cell specification and morphogenesis by removing bFGF and adding retinoic acid (RA). With this, spheroids underwent morphogenesis resembling neural tube formation. Such morphological conversions were easily visible under a transmission light microscope (Fig. 1b and Supplementary Video 2).

**Fig. 1.**
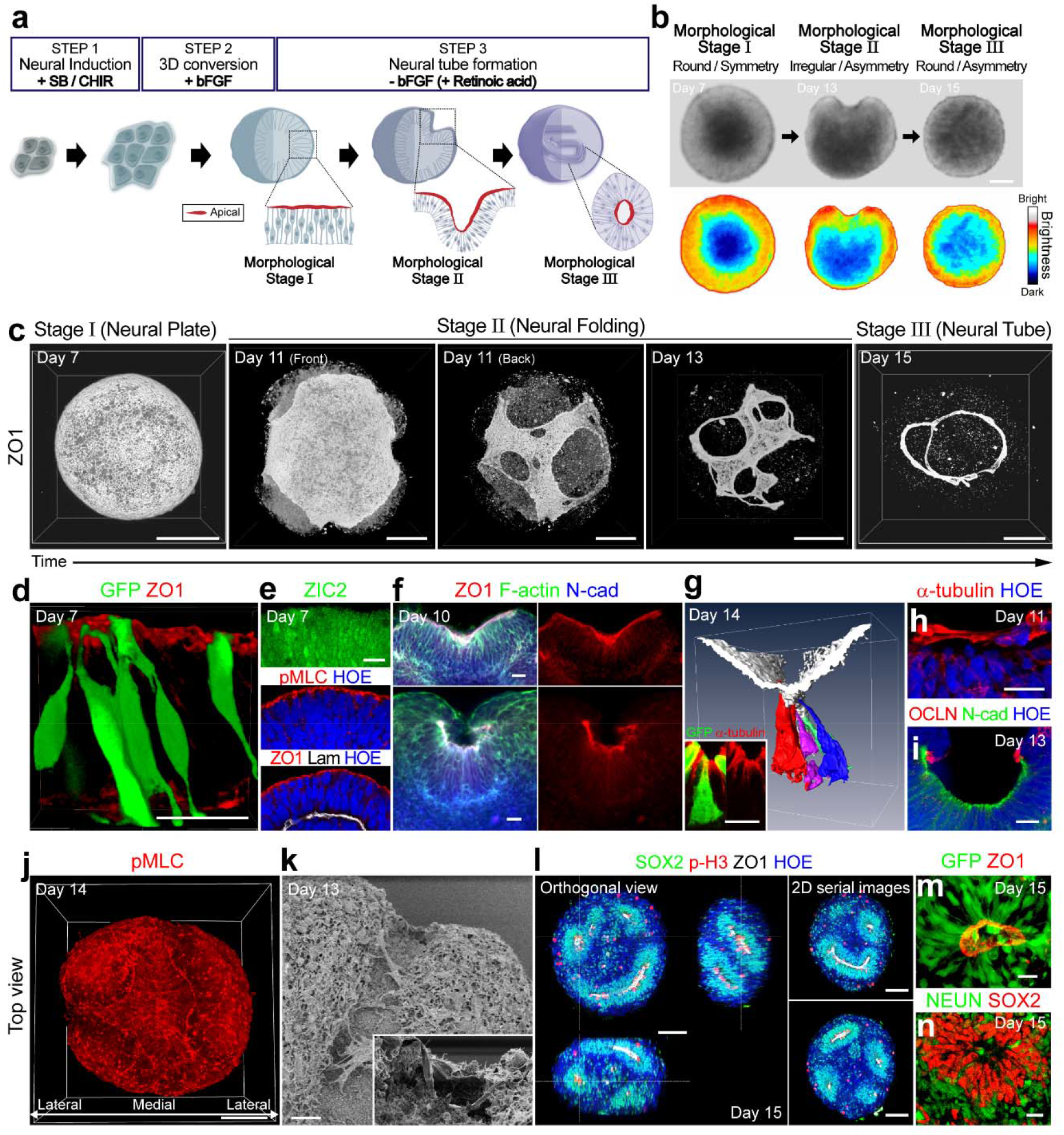
Generation of human spinal cord organoids (hSCOs) recapitulating neural tube morphogenesis. **a.** Schematics of the generation of hSCOs. **b.** Brightfield (upper) and pseudocolor (bottom) images at different stages of hSCO development. Scale bar, 100 μm. **c.** Time-course images of neurulation-like morphogenesis in hSCOs visualized with ZO1 (white) via 3D whole-mount imaging. Scale bar, 100 μm. **d.** Single-cell morphology of the NE layer. A small proportion (5%) of GFP-labeled H9 hESCs were blended with naïve H9 cells to better visualize the morphology of the individual cells. The apical side was labeled with ZO1 (red). Scale bar, 20 μm. **e.** Polarization of the NE cells was visualized with staining for ZIC2 (green), pMLC (red), ZO1 (red), and laminin (white). Nuclei were counterstained with Hoechst (blue). Scale bar, 20 μm. **f.** Two examples of different modes of neural folding. The upper image shows the hinge formation, and the lower image shows a simple round-up of neural plates. NE was visualized with ZO1 (red), F-actin (green), and N-cad (blue). Scale bar, 20 μm. **g.** Morphology of hinge cells in the hSCOs. Individual-cell morphology was visualized via GFP-labeled H9 cells, the apical side was labeled with ZO1 (white), and 3D images were processed with Amira software. The inset shows GFP+ cells with α-tubulin labeling (red). Scale bar, 20 μm. **h.** The cytosolic bridge is labeled with α-tubulin (red) covering the neural fold. Nuclei were counterstained with Hoechst (HOE, blue). Scale bar, 20 μm. **i.** Localization of OCCLUDIN (OCLN, red) at the dorsal tip of the neural fold. The apical side of the neural fold was labeled with N-cad (green). Scale bar, 20μm. **j.** Bird eye’s view of planar cell polarity of the NE cells visualized via pMLC during fold formation. Scale bar, 20 μm. **k.** SEM images of neural folding-stage hSCOs. The inset shows the position where the neural groove transformed into the neural tube. Scale bar, 20 μm. **l.** 3D image of neural tube-stage hSCOs on day 15 of culture. The tube structure was visualized with SOX2 (green) and ZO1 (white) staining. The mitotic cells expressing phosphohistone H3 (p-H3, red) were localized on the apical side of the neural tube. Scale bar, 100 μm. **m.** Single-cell morphology of NE cells in the neural tube. Scale bar, 20 μm. **n.** Double immunostaining for the neural stem cell marker SOX2 (red) and the neuronal marker NEUN (green). Scale bar, 20 μm.

Using the high-resolution 3D volume imaging method based on the tissue-clearing technique, we visualized the key characteristics of the organoid at each stage (Fig. 1c and Supplementary Video 3). In the presence of bFGF in step 2, the organoids established a distinct, columnar morphology of the NE surface layer exhibiting apical polarity (morphological stage I). Tracking the morphology of individual cells demonstrated a surface layer with an elongated pseudostratified columnar architecture (Fig. 1d). These cells exhibited the NE marker ZIC2^29^ and established apical polarity similar to the embryonic neural plate, such as apical localization of phospho-Myosin Light Chain (pMLC) (Fig. 1e). While the partial establishment of basal lamina in some organoids was observed, the basal lamina formation may not be essential for further morphogenesis as this feature was not prominent in certain organoids that underwent proper morphogenesis.

When the medium was replaced with -bFGF/+RA condition as a last step, round and symmetric spheroids began to fold (morphological stage II) (Fig. 1c, f, and Supplementary Fig. 2a). This process exhibited some features of neural folding *in vivo* in the perspective of morphogenesis. Along the apical side with an accumulation of cell-cell junctional proteins, the neural plate is gradually elevated and generate neural groove-like structures. At the cellular levels, the middle point cells in the neural groove showed the wedge-like cell morphology with condensed tubulin at the apical side (Fig. 1g and Supplementary Video 4). However, these wedge-like cells failed to express floor plate marker FOXA2 (Supplementary Fig. 2b), suggesting that the cellular differentiation and morphological alteration at the hinge points can be segregated. Although such NE layers in the organoid were not associated with any nonneuronal cells (Supplementary Fig. 2c), they exhibited a cytosolic bridge and strong clustering of occludin (Fig. 1h, *i)^30^.* The pMLC distributed laterally with planar cell polarity (PCP), producing cellular forces promoting neural folding, as noted *in vivo^31^* (Fig. 1j and Supplementary Fig. 2d-f). Scanning electron microscope images revealed the clear morphological change was observed at each stage in the organoid, especially at neural fold stage (morphological stage II), showing the continuum of groove-like neural folds and internalizing neural tube (Fig. 1k and Supplementary Fig. 2g). When tube morphogenesis was completed, the internalized neural tube exhibited an elongated tubular morphology, which was visualized by the apical localization of ZO-1 and pH3-expressing mitotic cells (morphological stage III) (Fig. 1l and Supplementary Video 5). They exhibited radial alignment of cells with apical polarity (Fig. 1m) and neural differentiation patterning by NEUN (Fig. 1n). The importance of cell polarity was further explored. The perturbation of cell polarization by the Rock inhibitor Y-27632 blocked apical polarity, and suppressed tube-forming morphogenesis (Supplementary Fig. 3a-c). Embedding the 3D spheroids into the matrigel rapidly inversed the polarity of the NE cells, and promoted morphogenesis resembling the rosette expansion-based ventricle formation as previously reported in brain organoid protocol^14, 18, 19^ (Supplementary Fig. 3d-f). We also verified this protocol with 4 different hPSC lines and consistently observed similar morphogenesis (Supplementary Fig. 4).

### Transcriptome analyses of hSCO maturation

Transcriptomic analysis was performed to evaluate the progress of hSCO maturation after initial morphogenesis period. The microarray dataset demonstrated a progress increase similarity in the global gene expression profiles between hSCOs and human fetal spinal cord tissue (Fig. 2a). Gene Ontology (GO) analysis demonstrated that neural stem cell proliferation and patterning-related genes were mainly expressed in the early stage hSCOS, while maturation-related gene clusters such as neurogenesis and gliogenesis were upregulated in late stage (Fig. 2b). To better understand the cell type specification of hSCOs at the single cell level, droplet-based single-cell RNA sequencing of 1-month hSCOs was performed with 11,038 cells. The clustering analysis was performed based on the representative transcription factors that identify the dorsal/ventral subclass cell types in spinal cord. The hSCOs included both mitotic and post-mitotic cells composed of approximate dorsoventral identity, mainly with the V0 domain cells (Fig. 2c, d and Supplementary Fig. 5a). hSCOs exhibited posterior identity indicated by HOX code associated with the cervical-thoracic level supporting their spinal cord identity (Supplementary Fig. 5b). While the domain-specific marker staining confirmed the presence of these cells in the hSCOs (Supplementary Fig. 6a), their regional patterning was not evident (Supplementary Fig. 6c). However, by treatments with dorsal inducer BMP4 or ventral inducer sonic hedgehog (SHH) agonist Purmorphamine hSCOs accordingly switched toward their domain identity, suggesting that they maintain the potential to form dorsoventral specification in response to the external morphogen stimuli. Thus, upon dissection of chick notochord as the source of ventral inducer and the treatment of recombinant BMP4, dorsoventral patterning was successfully established as observed by the position of chick notochord (Supplementary Fig. 6b-d). Re-clustering analysis according to the neurotransmitters identified 5 major clusters, namely, neural progenitors (SOX2), glutamatergic neurons (SLC17A6), glutamatergic/cholinergic neurons (SLC18A3), GABAergic neurons (GAD1), GABA/glycinergic neurons (SLC6A5), and other less-defined neuroblasts (DCX) (Fig. 2e-g, Supplementary Fig. 7).

**Fig. 2.**
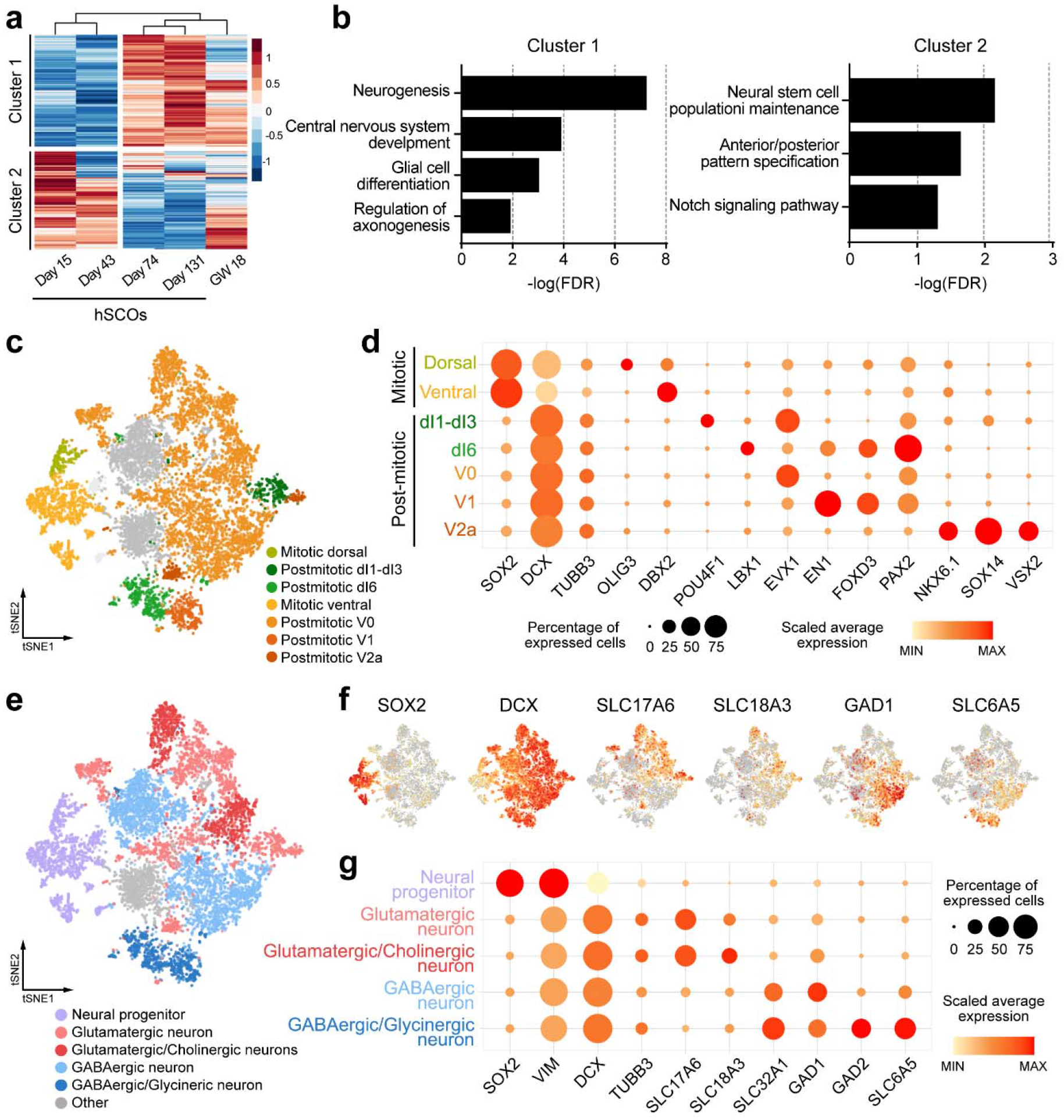
Transcriptome profiling of hSCOs. **a.** Heat map analysis of microarray data showing hierarchical clustering of 1556 differentially expressed genes identified from hSCOs and human fetal spinal cord tissue (gestational weeks 18). **b.** Significantly enriched GO terms related to biological process for upregulated (cluster 1) and downregulated (cluster 2) genes. **c.** Two-dimensional tSNE plot of 11,038 cells from 1month hSCOs by single-cell RNA-sequencing identified by clustering dorsoventral specific cells types. **d.** Gene expression profiles of the dorsoventral specific markers. Circle size and color represent the percentage of expressed cells and the average of gene expression, respectively. **e.** tSNE plot of re-clustered neuronal cell types from (c) datasets. **f.** The distribution of cells expressing the representative neuronal cell marker genes across the main populations. **g.** Gene expression profiles of the neuronal cell markers. Circle size and color represent the percentage of expressed cells and the average of gene expression, respectively.

### Morphological and cellular maturation of hSCOs

In the long-term culture, hSCOs grew and formed histological features of nerve-enriched shell and the cell-enriched core regions (Fig. 3a, Supplementary Fig. 8a and Supplementary Video 6). Consistent with the CNS development procedure, gliogenesis appeared to occur later, as early glial progenitor first seen by the 1-month and mature glial markers evident by 2-month (Fig. 3a,b and Supplementary Fig. 8b). Myelinating oligodendrocytes were also observed in mature organoids (Fig. 2c, d and Supplementary Fig. 8c).

**Fig. 3.**
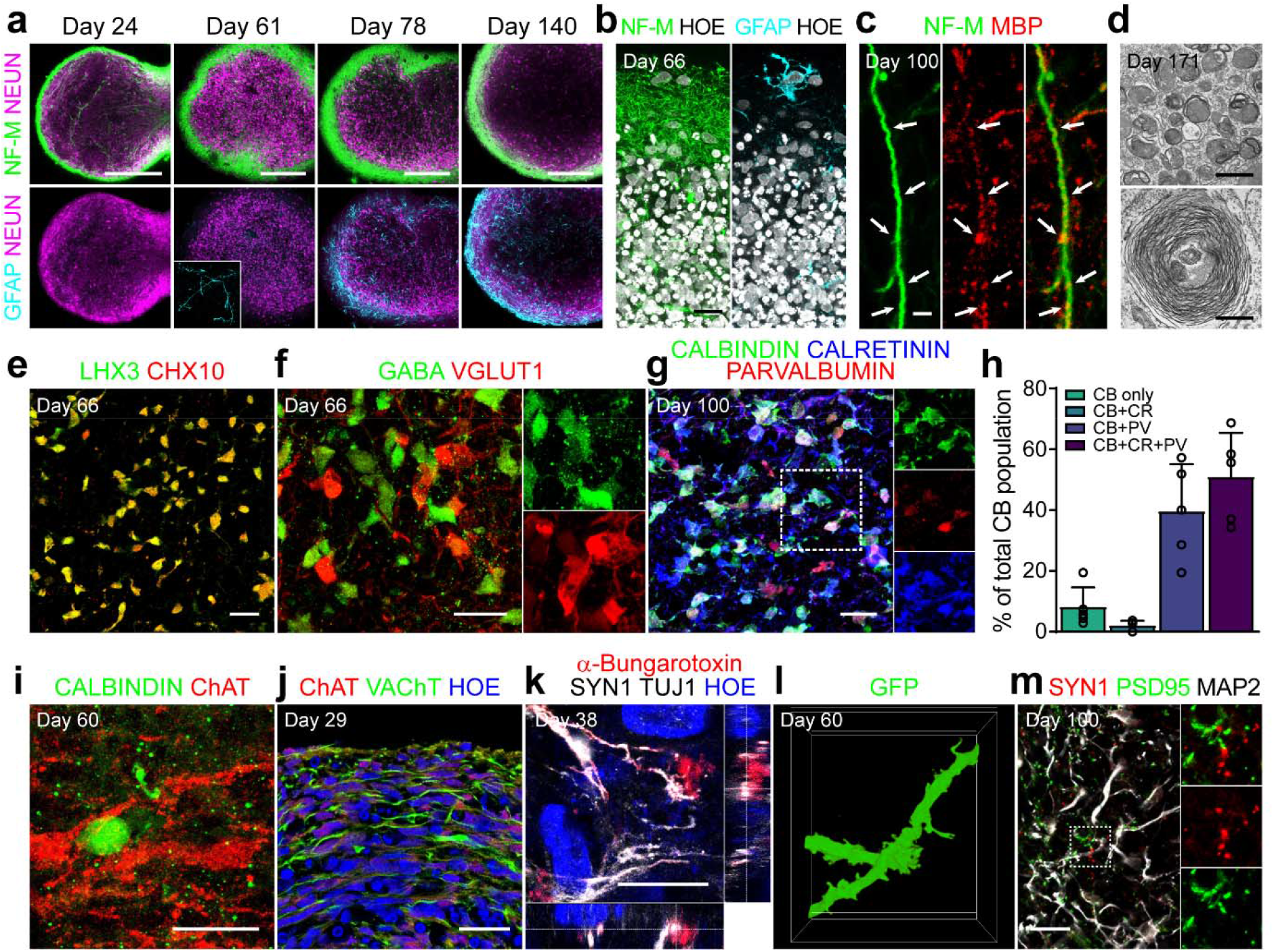
Acquisition of spinal cord–like cell fate after long-term culture of hSCOs. **a.** Maturation of hSCOs with structural segregation on the surface with neurite bundles and central neuronal cell bodies. The samples were stained with NeuN (magenta), Neurofilament-M (green), and GFAP (cyan). The inset shows the morphology of GFAP-positive astrocytes. Scale bar, 200 μm. **b.** High-magnification images on the surface. The samples were stained with Neurofilament-M (green) and GFAP (cyan). Nuclei were counterstained with Hoechst (white). **c.** Presence of oligodendrocytes labeled with MBP (red). White arrow indicates a neurofilament-M-labeled fiber (green) that was closely associated with an oligodendrocyte. Scale bar, 2 μm. **d.** Transmission electron microscopy images for the bundles of nerve fibers and myelination in hSCOs. Scale bar, 500 nm. **e.** Double immunofluorescent labeling of LHX3 (green) and CHX 10 (red) on day 66. **f.** Immunostaining for GABAergic (GABA, green) and Glutamatergic neurons (vGLUT1, rad) on day 66. **g.** Co-expression of CALBINDIN (CB, green), CALRETININ (CR, blue), and PARVALBUMIN (PV, red) on day 100. The inset shows triple-positive interneurons. **h.** Quantification of CB, CR, and/or PV co-expressing interneurons (error bars indicate s.e.m. n=5). **i.** Presence of Calbindin-expressing Renshaw cells (green) in proximity to ChAT-expressing motor neurons (red). **j.** Cholinergic neuronal axons running on the surface, visualized with staining for ChAT (red) and VAChT (green). **k.** Establishment of neuromuscular junction of outgrowing motor axons in human myotube coculture. **l.** The emergence of dendritic spine-like protrusions in the mature neurons of hSCOs. Morphology of the dendrites was traced from GFP-expressing neurons. **m.** Presence of mature synaptic markers, PSD95 (green), SYN1 (red), and MAP2 (blue) in the hSCOs. The inset shows a high-magnification image of the boxed area. All scale bars, 20 μm, except panel a, c, and d.

After >2 months, hSCOs exhibited V2a glutamatergic interneurons co-expressing LHX3 and CHX10, known as the reticulospinal neurons for locomotor functions^32^ (Fig. 3e). As shown in Fig. 2, both GABAergic (GABA+) and glutamatergic (vGLUT1+) interneurons were found with adjacent localization (Fig. 3f). Consistent with the specification of the GABAergic subtype during the development process *in vivo*^33^, Calbindin+ interneurons acquired heterogeneity, co-expressing other calcium-binding proteins such as calretinin and/or parvalbumin (Fig. 3g-h). Calbindin-expressing interneurons (Renshaw cells) formed contacts with nearby motor neurons—a typical neural network in the spinal cord for the reciprocal inhibition of motor neurons for pattern generation of alternate movements^34^ (Fig. 3i). Cholinergic neuron fibers (ChAT and VAChT) demonstrated a tendency to run on the surface of the hSCOs (Fig. 3j). When the hSCOs were co-cultured with differentiated human skeletal muscle myotubes, the outgrowing motor fibers had the potential to form neuromuscular junctions (NMJs) observed by labeling with α-bungarotoxin (α-Btx; Fig. 3k and Supplementary Fig. 9a-b). When hSCOs were co-cultured with dorsal root ganglions (DRGs) derived from Tau-GFP transgenic mice, sensory fibers from DRG were observed to readily penetrate the organoids, indicating that hSCOs can receive peripheral sensory input (Supplementary Fig. 9c-e). The neurons in the mature hSCOs exhibited a mature dendritic morphology with spine formation (Fig. 3l) and the expression of mature neuronal markers such as PSD95 and SYN1 (Fig. 3m) providing morphological evidence of synaptic contacts. In summary, our hSCOs demonstrated a transcriptional and histological similarity with the spinal cord *in vivo*.

### Spontaneous and evoked neuronal activity in spinal cord organoids

To evaluate the functional properties of mature hSCOs, we utilized the we utilized a MEMS neural probe embedded with microfluidic channels^35^. A single silicon neural probe with 16 microelectrode arrays was inserted into the middle of hSCOs (Fig. 4a), and the spontaneous neural activity was readily detected (an example of the traces was shown in Fig. 4b). Over time, the spontaneous neural firing/bursting rates increased progressively with highly synchronized neural activity—a typical feature of the developing spinal cord^36, 37^ (Fig. 4c-g and Supplementary Fig. 10a-l). It is noteworthy that burst activities increased rapidly as of 60 days when GFAP+ mature astrocytes were observed in hSCOs (Fig. 4e-f). Astrocytes are involved in the modulation of neural networks by regulating synaptic transmission as well as supporting neurons^38^. Therefore, after the advent of GFAP+ mature astrocytes in hSCOs, the rapid increase in burst activity is evidence that a complex functional neural network developed as hSCOs matured. Also, signal synchronization between the electrodes increased as the neural network expanded (Fig. 4g). After measurements of spontaneous neural activity under basal condition, we focally infused drugs during the recording via an outlet located in the silicon neural probe with a drug delivery channel for chemical modulation. Application of the voltage-sensitive sodium channel blocker tetrodotoxin (TTX) significantly suppressed the firing rate, thereby confirming that the electrical signals were a result of neuronal action potentials (Fig. 4h). Treatments with the AMPA receptor antagonist 6-cyano-7-nitroquinoxaline-2,3-dione (CNQX) and the NMDA receptor antagonist (2R)-amino-5-phosphonovaleric acid (AP5) significantly decreased the neural firing rate (Fig. 4i), confirming the excitatory synaptic transmission in hSCO. Treatment with GABA receptor antagonist bicuculline significantly increased the firing rate (Fig. 4j), while GABA receptor agonist baclofen decreased the neural firing rate (Fig. 4k), confirming the inhibitory synaptic transmission in hSCOs. Moreover, the infusion of Calcitonin gene-related peptide (CGRP) that activates spinal dorsal sensory interneurons significantly increased the neural firing rate, suggesting the functional maturation of sensory interneurons within hSCOs (Fig. 4l).

**Fig. 4.**
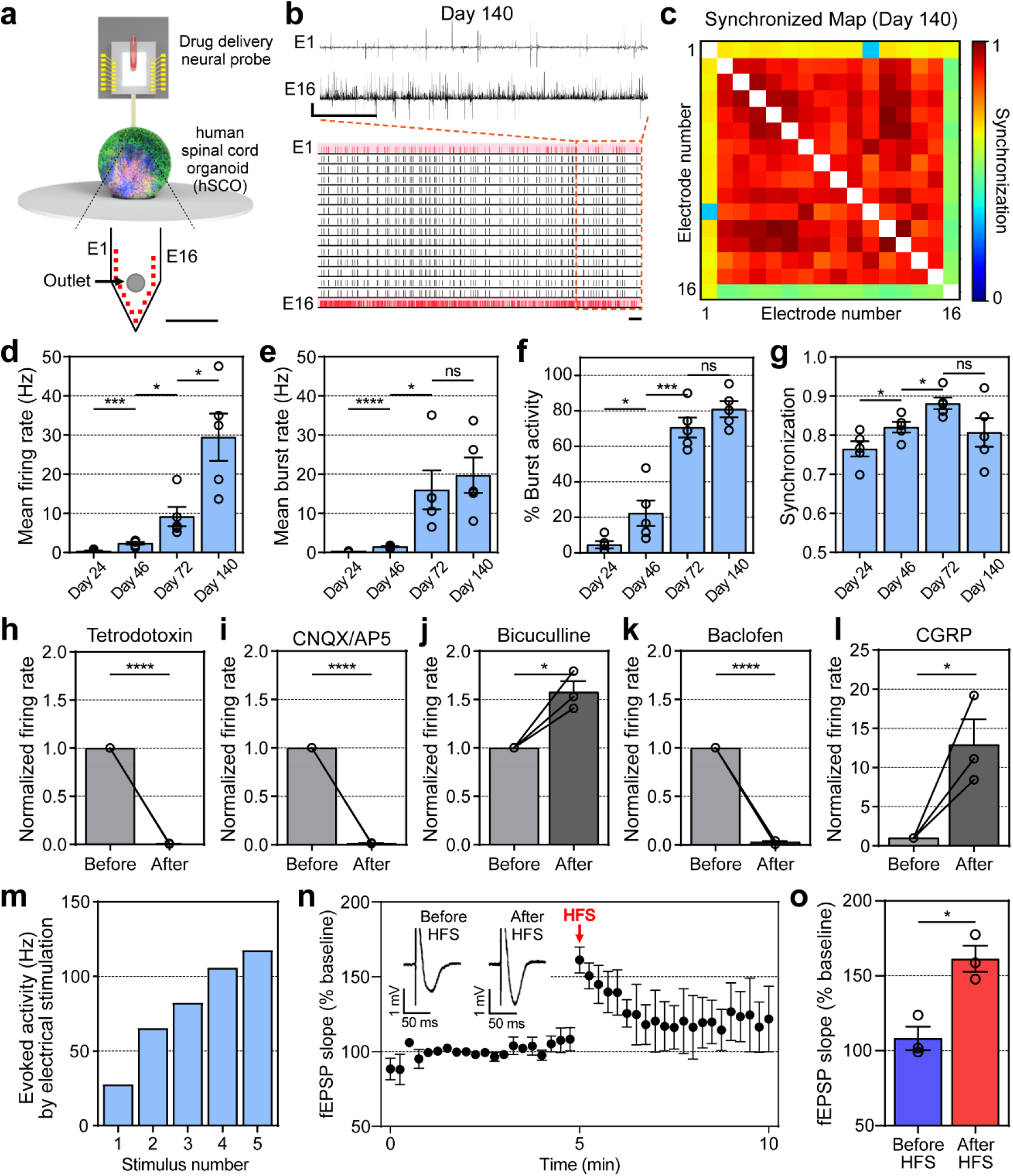
Electrophysiological analysis and pharmacological response in hSCOsa. Schematics of extracellular recordings from hSCOs using neural probe with drug delivery capability. Scale bar, 250 μm **b.** The representative raster and transient plot from neural activities recorded in hSCOs at Day 140. Scale bar, 1 sec (horizontal), 1 mV (vertical). **c.** The representative cross-correlation matrices showing synchronization between signal-recorded electrodes from hSCOs at Day 140. **d-g.** Changes in the patterns of neural activity upon hSCO maturation (error bars indicate s.e.m. n=5). Mean firing rate (Day 24 – 46: *** P 0.0007; Day46 – 72: * P 0.0244; Day 72 – 140: * P 0.0146; n=number of the samples, Two-tailed unpaired *t* – test) **(d).** Mean burst rate (Day 24 – 46: **** P < 0.0001; Day 46 – 72: * P 0.0192; Day 72 – 140: ns P 0.5960; n=number of the samples, Two-tailed unpaired *t*-test) **(e).** The percentage of burst activity in total activity (Day 24 – 46: * P 0.0445; Day 46 – 72: *** P 0.007; Day 72 – 140: ns P 0.1894; n=number of the samples, Two-tailed unpaired *t*-test) **(f).** Synchronization between electrodes (Day 24 – 46: * P 0.0467; Day 46 – 72: * P 0.0166; Day 72 – 140: ns P 0.0974; n=number of the samples, Two-tailed unpaired *t*-test) **(g). h-l.** Bar plots showing changes in the mean firing rate before and after drug treatments. 6 μM TTX (before – after: * P 0.0166; n=number of the samples, Two-tailed unpaired *t*-test) **(h).** 100 μM CNQX and 100 μM AP5 (before – after: * P 0.0469; n=number of the samples, Two-tailed unpaired *t*-test) **(i).** 10 μM Bicuculline (before – after: * P 0.0285; n=number of the samples, Two-tailed unpaired *t*-test) **(j).** 100 μM Baclofen (before – after: * P 0.0301; n= number of the samples, Two-tailed unpaired *t*-test) **(k).** 1 μM α-CGRP (before-after: * P 0.0333; n=number of the samples, Two-tailed unpaired *t*-test, error bars indicated s.e.m. n=3 for all) **(l). m.** Firing rate of electrically evoked activities according to stimulus number (The higher the repetition of the stimulus, the higher the increase in firing rate); **n.** Short-term potentiation in the matured hSCOs (error bars indicated s.e.m. n=3 for all; n=number of the sample). **o.** Bar plot showing the comparison of the fEPSP slope right before and after HFS (Before HFS-After HFS: * P 0.0105; n=number of the samples, Two-tailed unpaired *t*-test, error bars indicated s.e.m. n=3 for all);

Finally, we examined whether hSCOs exhibited neuronal plasticity, which is defined as the ability of the neurons to change network responses to previous intrinsic or extrinsic stimulations^39^. In 2-month hSCOs, the electrically-evoked activity showed a gradual increase in the action potential based on the repetition of electrical stimulations (Fig. 4m and Supplementary Fig. 10m). Short-term plasticity (STP) was evident as shown by the increase of field excitatory postsynaptic potential (fEPSP) slopes after high-frequency stimulation (HFS) (Fig. 4n and o). Taken together, these data suggest that the ‘learnable’ neural networks are well established with excitatory and inhibitory neural circuits within the hSCOs.

### Quantification of the effects of antiepileptic drugs (AEDs) on the tube morphogenesis in hSCO model

Finally, we tested whether the hSCO model can be used for drug screening or as a toxicology test platform. To improve the reproducibility of the organoid and make it suitable for the quantifiable assay of the tube morphogenesis, we optimized the procedure for 96-well plate-based high content analysis. After the initial induction of cNSCs in 2D, cells were dissociated into single cells, and a defined number of cells were seeded (Fig. 5a). The dissociated cells rapidly re-aggregated and re-established the outer NE layer, which is recognizable by the distinct nuclear alignments (Fig. 5b). As expected, the diameter increased with the cell seeding number, but NE thickness was not affected by the cell seeding number (Fig. 5c). In all sizes, the folding morphogenesis initiated similarly, but larger hSCOs spent longer period to complete morphogenesis (Fig. 5d-f and Supplementary Fig. 11a). Next, we evaluated the effects of the duration of bFGF treatment on the morphogenesis (Fig. 5g). By increasing the duration of bFGF treatments, both hSCO size and NE thickness were increased (Fig. 5h and i). hSCOs with a thicker NE layer spent longer time to initiate neural folding, and often failed to complete tube morphogenesis (Fig. 5j-l and Supplementary Fig. 11b). These data indicate that the kinetics of tube morphogenesis are tunable by modulation of initial cell number and the duration of bFGF treatment.

**Fig. 5.**
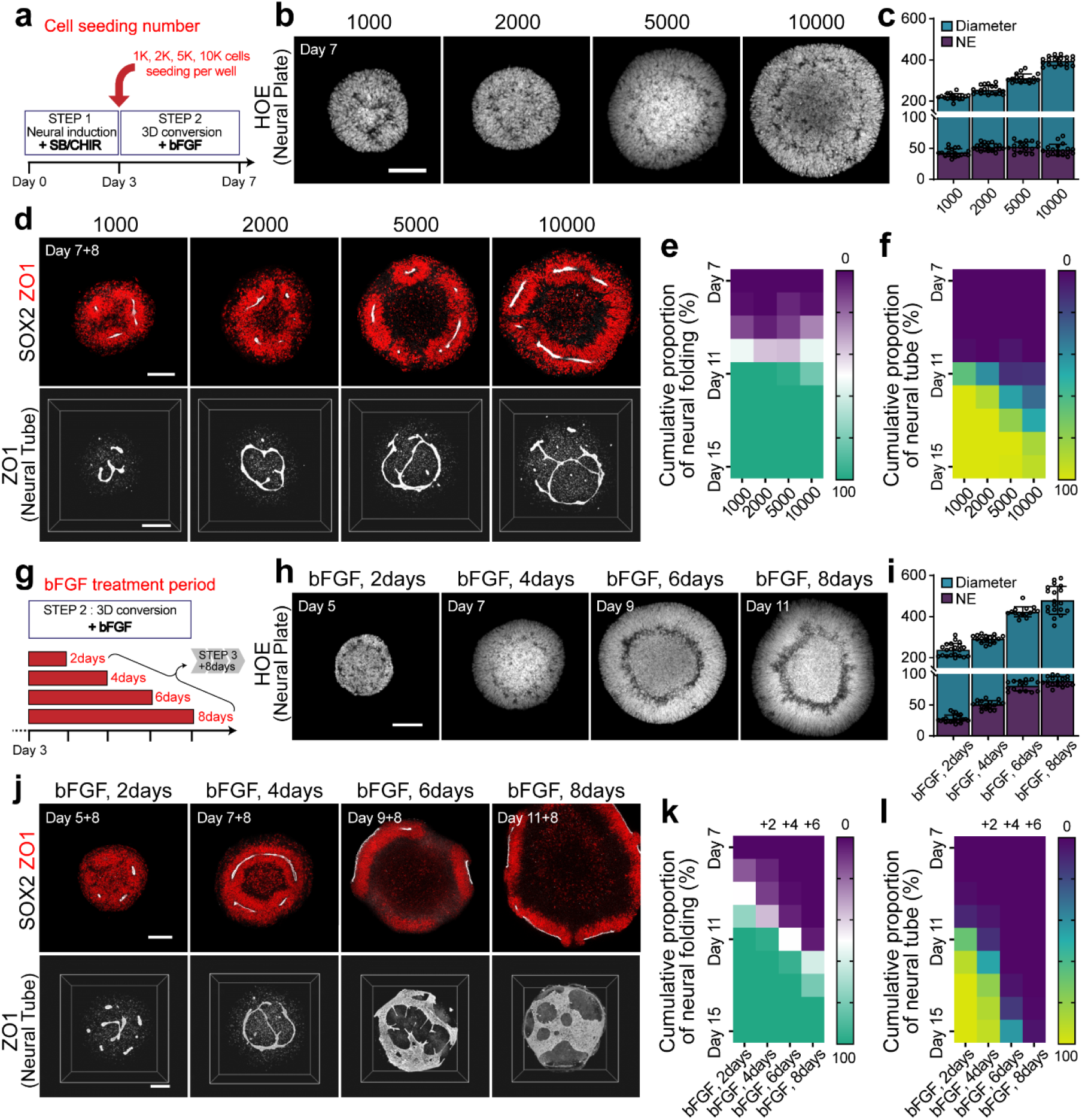
Size and bFGF duration-dependent tube morphogenesis in hSCOs. **a.** The experiment scheme to study the effect of different cell seeding numbers on morphogenesis. **b.** Neural plate formation with different initial cell seeding density. Nuclei were stained with Hoechst (white). **c.** Quantification of the diameter of hSCOs and NE thickness depending on the initial cell seeding numbers. **d.** Neural tube morphology visualized with SOX2 (red) and ZO1 (white) staining on day15. **e-f.** Quantification of morphogenesis with different cell seeding densities. The color of each box indicates the cumulative proportion of the neural folding (**e**) or neural tube (**f**) stage of hSCOs at the indicated culture time. **g.** The experiment scheme to examine the effect of bFGF treatment durations. **h.** Neural plate formation with different bFGF treatment durations. Nuclei were stained with Hoechst (white). **i.** Quantification of the diameter of individual hSCOs and NE thickness depending on the duration of bFGF treatments. **j.** Neural plate/tube morphology visualized with SOX2 (red) and ZO1 (white) staining. **k-l.** Quantification of tube morphogenesis with increasing duration of bFGF treatments. The color of each box indicates the proportion of cumulative neural folding **(k)** or neural tube **(l)** stage of hSCOs at the indicated culture intervals. All scale bars, 100 μm.

As the morphogenetic process can be imaged, we employed deep learning-based image analysis tools as a cost-effective and automated analysis system. For the establishment of an automated analysis system, we prepared a “pre-labeled” dataset comprising approximately 2000 organoids and trained an algorithm with supervised learning to automatically classify the stage of morphogenesis (Fig. 6a). In summary, this framework includes 1) An automatic imaging system that tracks the individual morphogenesis of each organoid in 96-well plates, 2) Deep learning-based classification of the morphogenetic process, and 3) Volume imaging analysis of neural tube structure.

**Fig. 6.**
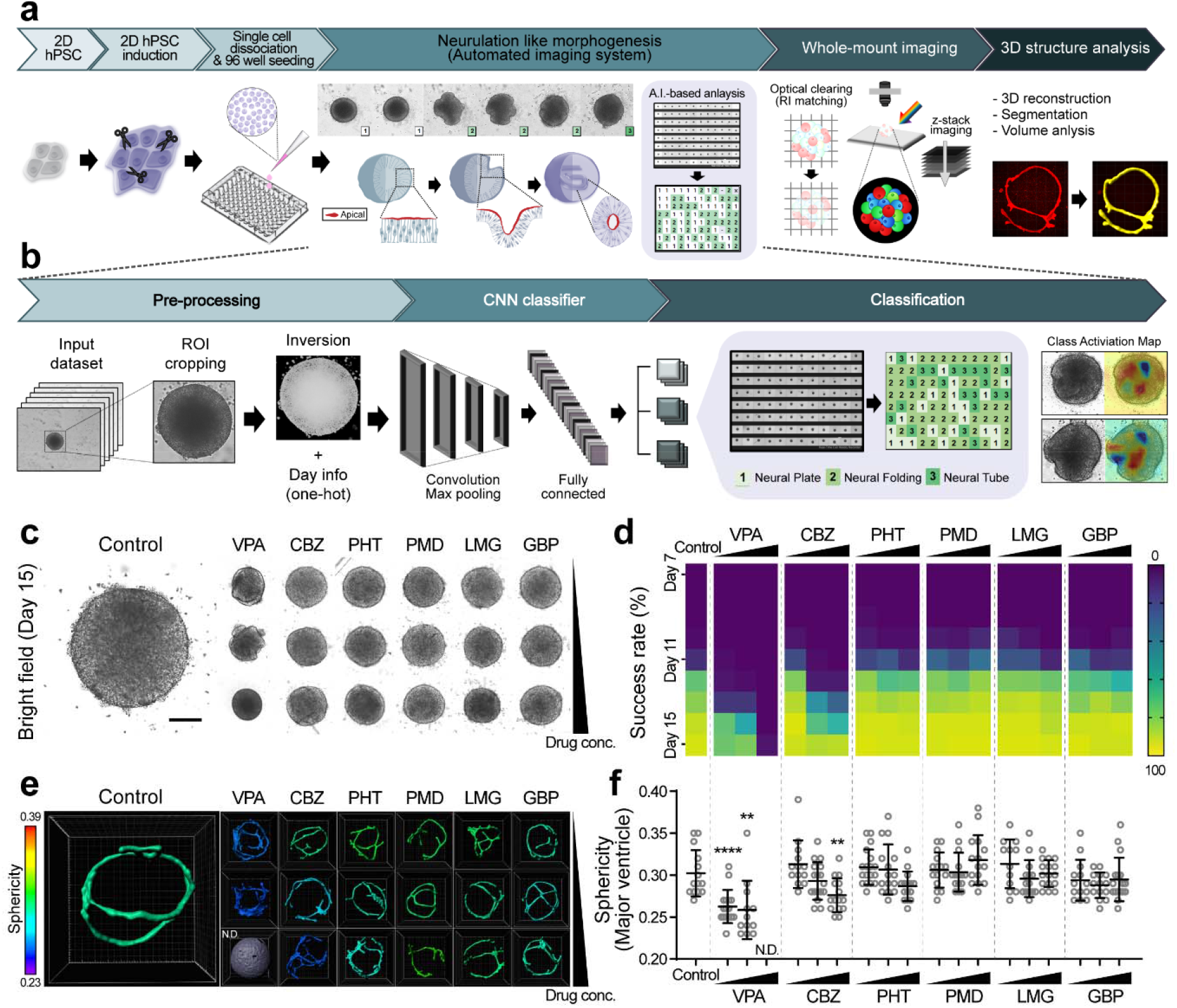
Modeling of antiepileptic drug-induced NTDs in hSCOs using deep learning-based classification. **a.** The workflow of an optimized protocol for the quantitative analysis of the neurulation-like process. After re-seeding the dissociated cells into a microwell (96-well) plate, the morphology of each organoid was captured with a 1-day interval using an automated imaging system. **b.** Training procedure for a supervised machine learning model. Based on the bright field images acquired in panel (a), the morphogenic stages (I: Neural Plate, II: Neural Folding, III: Neural Tube) were scored and further analyzed. **c.** Bright-field images of drug-treated hSCOs on day 15. Scale bar, 100 μm. **d.** Quantification of morphogenesis after treatments with six different AEDs. Each box shows the proportion of hSCOs exhibiting the neural tube. **e.** 3D neural tube morphology was labeled with ZO1 staining after treatments with six different AEDs. Pseudocolor is based on the sphericity value, which is calculated by the volume and surface area. **f.** Scatter dot plots for the individual sphericity of 3D neural tube measured by ZO1-labeled lumens. (Control – VPA low dose: **** P < 0.0001, n=16; Control – VPA middle dose: ** P = 0.0015, n=12; Control – CBZ high dose: ** P = 0.0069, n=15; n=number of the samples, Two-tailed unpaired *t*-test)

To test the proof-of-concept of the system, our platform was analyzed for efficient detection of the effects of AEDs on the NTDs. NTDs are common congenital malformations with a frequency of 0.5–2 per 1,000 pregnancies^40^. Open NTDs, such as spina bifida, are caused by the failure of neural tube closure, which can be a result of genetic mutations or environmental factors^5, 41^. Especially, AEDs increase the risk of the development of NTDs in humans^9, 42^. To validate our approach, hSCOs were treated with three different concentrations of 6 selected AEDs, including valproic acid (VPA), carbamazepine (CBZ), phenytoin (PHT), primidone (PMD), lamotrigine (LMG), and gabapentin (GBP). On day 15 following the 8-day treatment of AEDs, VPA- and CBZ-treated groups exhibited a failure of neural tube closure in a dosedependent manner, while others exhibited normal morphogenesis (Fig. 6c). The heatmap images based on the proportion of NT showed that the initiation of morphogenesis was retarded by the VPA or CBZ treatments, resulting in the delay or failure of tube closure (Fig. 6d and Supplementary table). High-resolution 3D imaging with ZO1 as an apical neural tube marker confirmed that the NTD-like phenotypes developed in response to VPA or CBZ treatments such as incomplete closure and increased branching of the internalized tube structures (Fig. 6e, f and Supplementary Video 7). These results were highly consistent with the reported clinical risks of AEDs, in which VPA and CBZ exhibited the highest risk ratio of congenital malformation, primarily NTDs^42^. Thus, the hSCOs-based toxicology system can recognize the effects of drugs on early neural development, such as the development of NTDs.

## Discussion

Here, we established a robust and quantifiable method for the production of hSCOs via neurulation-like morphological processes. While organoids representing a specific part of the CNS have been reported, they grow via rosette^14–17^ or incomplete folding formation^19^ with subsequent expansion of ventricles, whose morphogenesis is similar to follicular enlargement. However, our protocol allowed the tube-like morphogenesis resembling the neural tube formation *in vivo.* The prolonged culture of the hSCOs drove the organoids toward the development of the spinal cord. They exhibited the emergence of the spinal cord-type neurons and glial cells, and synaptic network activity. While studies have reported spinal cord-like organoids or neural tubes^11, 20–24, 43–45^, no study has comprehensively examined the later maturation steps. Particularly with the employment of a nanoelectrode system, electrophysiological properties of hSCOs were easily detectable. We optimized the protocol for culturing individual organoids to quantifiably assess the morphogenesis process, and the robustness of the protocol was successfully demonstrated by the quantifiable examination of the effects of drugs on the tube morphogenesis. Since hSCOs further mature into spinal cord-like neural circuits, our current culture system will provide a valuable opportunity for large-scale analysis of the development and function of human spinal cords *in vitro.*

The neural tube is an embryonic neural tissue, which leads to the development of CNS^1^. It develops through an early developmental process called neurulation, a complex yet ordered process of neural cell specification and morphogenesis that includes early induction of the neural plate, neural fold formation, and neural tube closure. In early embryonic tissue *in vivo*, the neural plate is a planar structure connected to the surface ectoderm. In this model, the neural plate-like structures emerged as a surface layer of the entire 3D sphere. This neural plate-like layers exhibited proper apicobasal and planar cell polarities similar to the neural plate *in vivo*, which is essential for initiating tube morphogenesis that occurs by symmetry breaking. The hSCOs do not contain any non-neural tissue components, such as non-neural epithelia or mesodermal tissues. The core neurulation program is influenced by the surrounding non-neural tissues such as notochord and non-neural epithelia. However, very few studies have examined the contribution of NE cells and other surrounding tissues for neurulation, rendering the conclusion elusive^46–48^. NE layer underwent tube morphogenesis without the non-neural components in hSCOs. Therefore, it may represent a self-organizing process of NE. This finding is consistent with observations based on many organoids in which epithelial linings can form unique tissue architectures without mesenchymal support^49^.

Although hSCOs replicated some features of neural tube formation *in vivo,* several differences were observed between the hSCO morphogenesis and *in vivo* neural tube formation. For instance, while we observed wedge-like cells emerging at the base of the folding NE layer undergoing tube formation, they did not exhibit floor plate markers. In addition, *in vivo* neural tube formation is associated with border formation and the induction of neural crest progenitors. However, we failed to identify neural crest lineage induction. Also, hSCOs *per se* did not exhibit dorsoventral patterning, although they responded to the supplements with chick notochord and treatment with dorsal induction signal BMP4. We speculate that these differences are primarily caused by the absence of non-neural components. Thus, this organoid model will help clarify the self-organizing events of NE cells versus the modulatory role of non-neural tissues in neural tube morphogenesis.

In the aspect of quantification analysis, this hSCO protocol provides a unique advantage over other protocols generating neural organoids. Many neural organoids are induced from 3D spheroids, and the tissue-organizing centers appear to emerge randomly, resulting in complex repeats of follicular tissue architectures. Although some protocols for dorsal telencephalic organoids offer smaller variations in terms of size and cell compositions^17, 26, 50^, variations in the organoids developed in many protocols are one of the major problems for the application of organoids in a quantifiable assay system. We induced a neural fate in 2D and re-aggregated the dissociated cells with a defined number, leading the hSCOs to undergo uniform morphogenesis as a “unit”, suitable for the rapid, simple, and accurate quantifiable analyses. The advantage of this protocol includes the ability to produce a large quantity at a time, to control the size and speed of organoid development, and to automate the imaging system tracking individual morphogenesis, particularly suitable for applications such as high-throughput toxicological screening. The use of an AI-based analysis toolkit allowed unbiased drug screening and can be implemented with an automated system. The testing of our system with AEDs shows the potential and reliability of our hSCO model. Considering that NTD has no reliable toxicology tests available, our demonstration is a good example of how neural organoids can be used for drug screening and toxicology tests as new approach methodologies (NAMs).

## Methods

### Human PSC Culture

Human H9-GFP ESCs were modified from H9 ESCs by lentiviral mediated insertion of a GFP-expressing cassette into the cellular genome, followed by colony selection. Human iPSCs #5-1 were derived from epidermal fibroblasts, and #56-2 were derived from peripheral blood monocytes. Human iPSCs AICS-0023 were purchased from Coriell Institute. Human PSCs were maintained on matrigel (BD Biosciences, 354277)-coated plates in E8 medium (STEMCELL Technologies, 05990) for hESCs or mTeSR1 (STEMCELL Technologies, 85850) for hiPSCs. The hESCs and hiPSCs were maintained under 5% CO2 at 37°C with daily media change and were passaged every 5 to 7 days by ReLeSR (STEMCELL Technologies, 05872) into small clumps and replated onto the precoated culture dishes. Experiments were performed on hiPSCs below passage 50.

### Generation of human spinal cord organoids

To generate human spinal cord organoids (hSCOs), dissociated small clumps of hPSCs were plated onto matrigel-coated plates at high density in mTeSR1. After cell attachment, mTeSR1 was replaced with the differentiation medium (DM) consisting of DMEM/F-12. To promote the induction of caudal neural stem cells, hPSCs were treated with SB431542 and CHIR99021 in DM for 3 days with daily media change^27^. On day 3, intact colonies were gently detached from dish. Detached colonies were then transferred onto uncoated culture dishes in DM supplemented with basic fibroblast growth factor (bFGF). They began forming a neuroepithelial (NE) structure at the peripheral surface of the organoid and were fed daily for 4 days. On day 7, hSCOs were cultured in the DM containing retinoic acid (RA) without bFGF for 8 days, inducing neural plate morphogenesis in NEs to form the neural tube. The medium was changed every other day. To disrupt cell polarity, the hSCOs were cultured in DM containing Y27632 (10 μM, Tocris, 1254) for 4 days or embedded within a matrigel in DM for 3 days (following the formation of a spheroid on day 4) with daily media change. Brightfield images were acquired with an EVOS microscope (Life Technologies) to observe morphological alterations during neural plate morphogenesis. For maturation, hSCOs were grown in 1:1 mixture of DMEM/F-12 and neurobasal medium. The medium was changed every 3 to 5 days.

A standard protocol was followed using 96-microwell plates. On day 3 of differentiation, the attached cells were dissociated with Accutase (Innovative Cell Technologies, AT-104). To observe morphogenesis under optimal conditions, the dissociated cells were seeded onto a 96-well low attachment plate (5000 cells per well) in DM supplemented with 20 ng/mL bFGF. The hSCOs in the microplate were fed daily for 4 days. On day 7, hSCOs were cultured in DM containing retinoic acid (RA) without bFGF for 8 days. To induce neural tube defects by AEDs, the hSCOs were either cultured in DM containing AEDs (valproic acid, 0.5 or 1 or 2 mM, Sigma, P4543; carbamazepine, 5 or 50 or 100 μM, Sigma, C4024; phenytoin, 3 or 30 or 100 μM, Sigma, D4007; lamotrigine, 30 or 100 or 300 μM, Tocris, 1611; primidone, 5 or 50 or 150 μM, Tocris, 0830; gabapentin, 10 or 300 or 100 μM, Tocris, 0806). The medium was changed every day. Brightfield images were automatically acquired with JuLi™Stage (NanoEntek) to quantify the rate of neural plate morphogenesis.

### Tissue clearing and 3D imaging

For 3D volume imaging, hSCOs were fixed with 4% paraformaldehyde (PFA, Biosesang) for 30 min (< 500 μm diameter) or 1 h (> 500 μm), followed by washing several times with PBST (0.1% Triton X-100 in PBS), and incubating with a blocking solution (6% BSA, 0.2% Triton X-100, and 0.01% sodium azide in PBS) overnight. For 3D wholemount immunostaining, hSCOs were immersed in primary antibodies diluted in blocking solution for 48 h (Antibodies and their dilutions are listed in Table S1). The primary antibody was then washed with PBST thrice for 10 min. Subsequently, hSCOs were incubated with the appropriate secondary antibody and Hoechst33342 diluted in a blocking solution for 48 h. The organoids were then washed with PBST thrice for 10 min and mounted onto the coverglass (24 x 40 mm) with a mounting solution (25% urea and 65% sucrose in H2O) for optical clearing. All steps were performed in a 0.6 mL tube or 0.2 mL tube with gentle shaking at RT. All images were captured with a Leica TCS SP8 Confocal microscope.

### 3D Image processing and volumetric analysis

For 3D imaging and analysis of hSCOs, raw images were collected using Leica SP8 and processed with LAS X software (Leica). The 3D images of hSCOs were created using Z-stacks (typically 50–300 images) with 0.5–2 μm intervals, and then manually segmented and rendered with the AMIRA software (Thermo Fisher). The regions of interest (ROI) were manually defined based on the intensity of images. For volumetric analysis of neural tube, raw images were rendered with the IMARIS software (OXFORD Instruments). The segmentation of ROIs was performed by intensity-based thresholding with labeled ZO1. The 3D structural quantification including parameters such as sphericity, volume, and surface area were calculated by the IMARIS software. The sphericity (*Ψ; π*^1/3^(6*V*)^2/3^*A^-1^*) was calculated by the volume *(V)* of ZO1 particles and its surface area *(A).*

### Histology and immunofluorescence

The hSCOs were fixed by immersion in 4% PFA overnight at 4°C and washed several times in PBS. The samples were then incubated with 30% sucrose in PBS at 4°C until completely submersed, embedded in Tissue-Tek Optimal Cutting Temperature (O.C.T. Compound, SAKURA), frozen on dry ice, cryosectioned serially to obtain 16-to 40-μm thickness and collected onto New Silane IIWE coating slides (Muto Pure Chemicals Co. Ltd, 5118-20F). For immunostaining, samples were permeabilized with PBS thrice with 5 min durations at RT, blocked with a solution (3% BSA and 0.2% Triton X-100 in PBS) for 30 min at RT, and incubated with the respective primary antibody diluted in a blocking solution overnight at 4°C. (Antibodies and their dilutions are listed in Table S1.) Samples were then washed with PBS thrice for 5 min durations at RT and then incubated with the respective secondary antibody and Hoechst33342 diluted in the blocking solution for 30 min at RT. The secondary antibody was subsequently washed with PBS, and the samples were mounted in the Crystal mount (Biomeda, M02). All steps were performed with gentle shaking. Images were acquired and processed with the Leica TCS SP8 Confocal microscope system.

### RT-PCR analysis of gene expression

Total RNA was isolated from hSCOs using Trizol (Invitrogen) or the RNeasy Mini kit RNase-Free DNase set (QIAGEN) in triplicate according to the manufacturer’s instructions. Isolated RNA (1 μg) was used to synthesize cDNAs using Murine Moloney Leukemia Virus reverse transcriptase (MMLV, Promega). Subsequently, cDNAs were amplified with genespecific primers (Sequence of primers are listed in Table S2). PCR conditions and the number of cycles (25-35) were optimized as follows: 95°C for 15 min, denaturation at 95°C for 30 s, annealing at 58-60°C for 30 s, and extension at 72°C for 30 s. qRT-PCR (Applied Biosystems, ABI7500) analysis was performed using the SYBR GREEN master mix (Enzynomics or Elpis) in combination with specific primers. The reactions were performed with an Eppendorf Realplex4 cycler (Eppendorf). All values were normalized to GAPDH expression for calculating the fold change.

### Machine learning

The dataset was acquired on JuLi™Stage (NanoEntek) with a 4x objective lens. In preprocessing, raw images were cropped into 512×512 pixel windows except for the background. This step was automatically proceeded by the brightness distribution of the image. To minimized batch effects arising from the differences in brightness/contrast, the pixel values were normalized with even distribution. Of the total dataset comprising over 12,000 images, we used 90% for training and 10% for validation at each morphological stage. In the training set, image augmentation was used to increase the dataset size by randomly applying rotation, translocation, flipping, zooming in, and zooming out. Model design and training were performed using a convolutional neural network (CNN) in Tensorflow 1.13. The architecture of the CNN model consisted of 5 convolution layers, max-pooling, and 2 fully connected layers. To improve CNN classifier performance, the culture day-information was reflected as metadata by One-Hot encoding. Finally, the classification results of morphological stages were presented as 1 (neural tube), 2 (neural folding), or 3 (neural tube) as well as the class activation mapping (CAM) to visually validate images along with the prediction scores.

### Electrophysiology

To evaluate the functionality of the cultured hSCOs, we used a MEMS neural probe integrated with 16 Pt microelectrodes and a microfluidic channel for neural signal recording and localized drug delivery^51^. The following steps were performed: 1) To improve the recording and stimulation capabilities, black Pt-coated microelectrodes (19 μm x 19 μm) were employed^52^. 2) The impedance of black Pt-coated microelectrodes was measured in 0.1 M PBS with a saturated calomel electrode using an impedance analyzer (nanoZ, Neuralynx). The average impedance was 13±1 kΩ at 1 kHz. To monitor neural activities of the organoid in an incubator and inserting the neural probe into the organoid, we used a small-sized custom microdrive system. The microdrive system consisted of 1) a PDMS recording chamber for the loading of the organoid, 2) a microdrive for adjusting the vertical position of the neural probe, and 3) an acrylic box for preventing rapid media evaporation in the incubator. After fixing the neural probe on the microdrive using two screws (1 mm x 3 mm), we transferred the organoid into the recording chamber. After positioning the organoid under the neural probe, the sample was embedded in low-melt agarose, and then the neural probe was slowly inserted into the organoid via the microdrive. The recording chamber was filled with a fresh DMEM/F 12-based culture media. After placing the organoid with the neural probe inserted into the acrylic box, we measured the neural activities of the organoid in the incubator.

Signals recorded from 16 black Pt-coated microelectrodes were processed and digitized using an RHD2132 amplifier board connected to an RHD2000 Evaluation System (20 kS s-1 per channel, 300 Hz high pass filter, 6 kHz low pass filter, 16-bit ADC for spike recording, 20 kS s-1 per channel, 0.3 Hz high pass filter, 3 kHz low pass filter, 16-bit ADC for fEPSP recording). Spontaneous neural activity was recorded for at least 10 min and from at least 3 organoids. Additionally, to deliver drugs (TTX, CNQX/AP5, Bicuculine, Baclofen, CGRP) to the sites where neural activity was measured in hSCOs, we used embedded microfluidic channels in the neural probe. For a faster response time, a pressure-driven drug delivery system was used. An electro-pneumatic regulator (ITV0051-2BL, SMC Pneumatics, Tokyo, Japan) was connected to a nitrogen tank to control the precise input pressure. After monitoring spontaneous neural activities of the hSCOs, 1.5 μL of fresh medium was administered with the drug for 3 min at a flow rate of 500 nL/min, and the change in neural activity following the injection was monitored for 3 min. In this experiment, we used 6 μM TTX, 100 μM CNQX, 100 μM AP5, 10 μM bicuculline, 100 μM baclofen, and 1 μM α-CGRP. For measurements of the evoked activity by electrical stimulation and short-term plasticity (STP), A365 stimulus isolator (WPI, Sarasota, FL, USA) was used for electrical stimulation. Some of the microelectrodes on the neural probe were used as stimulating electrodes. Evoked activity and STP were induced by 1 train of HFS (20 pulses 100 Hz, 200 μs, 100 μA for evoked activity, 100 pulses 100 Hz, 200 μs, 100 μA for STP). In the case of STP measurement, neurons in the hSCOs were stimulated by single pulses (50 μA, 200 μs pulse width, 30 s inter-pulse interval) before and after HFS for fEPSP recording.

To detect neural activities from recorded signals, we used a custom Matlab spike-sorting algorithm^35^. The threshold amplitude (75 μV) was set at more than three times the noise level (~ 25 μV). Burst activities among the detected neural signals were analyzed using the ISIN-threshold method (ISI threshold: 0.1 sec, minimum number of spikes: 3)^53^. The synchronized activity between electrodes was analyzed using Pyspike (https://github.com/mariomulansky/PySpike).

### Scanning electron microscopy and transmission electron microscopy

Samples were fixed with 2.5% glutaraldehyde in 0.1 M phosphate buffer at 4°C for 2 h before being washed with 0.1 M phosphate buffer thrice at RT. Fixed hSCOs were then subjected to a secondary fixation procedure—they were soaked in 1% osmium tetroxide in 0.1 M phosphate buffer for 90 min at RT. Subsequently, hSCOs were dehydrated via a series of ethanol (60%, 70%, 80%, 90%, and 95%) washes for 15 min each, followed by three washes with absolute ethanol with the duration of 30 min at RT. Dehydrated hSCOs were immersed in tert-Butyl alcohol twice for 20 min each at RT. Subsequently, hSCOs were frozen at −70°C and freeze-dried to remove tert-Butyl alcohol. Finally, hSCOs were mounted on the top of a sample holder with carbon tape, coated with platinum, and viewed under a scanning electron microscope (Hitachwe S-4700, Hitachwe High-Technologies Corporation, Tokyo, Japan).

For transmission electron microscopy (TEM), hSCOs were pre-fixed with 2% PFA and 2.5% glutaraldehyde in 0.1M phosphate buffer (pH 7.4) at RT for 1 h. Fixed hSCOs were then washed twice with 0.1M phosphate buffer (pH 7.4) and post-fixed with 1% osmium tetroxide in 0.1M phosphate buffer (pH 7.4) for 1 h at RT. To increase the image contrast, en bloc staining was performed with 0.1% uranyl acetate in 50% ethanol for 1 h. Subsequently, the hSCOs were dehydrated via an ascending ethanol series, followed by embedding in Epon812 (Okenshoji, Japan) and polymerization in a dry oven (65°C, 48 h). Tissues were sectioned (70 nm) using an EM UC7 ultra-microtome (Leica), mounted onto 200-mesh copper grids, and stained with 2% uranyl acetate and lead citrate for 5 min each. Sections were observed by TEM (Hitachwe H-7650, Hitachwe High-Technologies Corporation, Tokyo, Japan).

### Single-cell RNA sequencing

The hSCOs were collected in a petri dish on day 29 and chopped into small pieces. After dissection, the hSCOs were dissociated by using papain containing L-cysteine by incubation at 37°C with gentle shaking. Papain was removed after 30 min, and the dissociated cells were washed twice with ice-cold HBSS. Libraries were prepared using the Chromium controller according to the 10x Single Cell 3’ v3 protocol (10x Genomics). Briefly, the dissociated cell suspensions were diluted in nuclease-free water to achieve a targeted cell count of 10,000. The cell suspension was mixed with a master mix and loaded with Single-Cell 3’ Gel Beads and Partitioning Oil into a Single Cell 3’ Chip. RNA transcripts from single cells were uniquely barcoded and reverse-transcribed within droplets. cDNA molecules were pooled, and the cDNA pool then went through an end repair process, followed by the addition of a single ‘A’ base and ligation of the adapters. The products are then purified and enriched with PCR to develop the final cDNA library. The purified libraries were quantified using qPCR based on the qPCR Quantification Protocol Guide (KAPA) and qualified using the Agilent Technologies 4200 TapeStation (Agilent technologies). The libraries were subsequently sequenced using the HiSeq platform (Illumina) with 33,000 reads/cell.

### Preprocessing and analysis of single-cell RNA-Seq data

The raw sequence data were processed using the Cell Ranger pipeline (version 3.1.0, 10x Genomics). Reads were aligned to the GRCh38 human genome reference using the STAR aligner (version 2.5.1b)^54^. Gene expression matrices were generated using the Seurat package (version 3.1.5)^55^ in R. Unless otherwise stated, we used functions of the Seurat package for downstream analysis. Several quality-control steps were performed to filter out unreliable cells and genes as follows: (i) removal of cells with >10 % of counts that were mapped to mitochondrial genes, (ii) removal of cells with more than 7,000 or fewer than 200 unique expressed genes, (iii) removal of genes detected in <10 cells. After the quality control procedure, we obtained 18,365 genes across 11,038 cells for further analyses.

For downstream analysis, the UMI count data were normalized and variance-stabilized using the R package SCTransform^56^. The effect of mitochondrial gene expression was removed during the normalization process. Highly variable genes were selected based on the variance of Pearson residuals from normalized negative binomial regression. A graph-based clustering algorithm (FindClusters function of Seurat) was used for single-cell clustering. The t-Stochastic Neighbor Embedding (tSNE) visualization was achieved using the top 20 principal components obtained from principal component analysis based on the elbow plot and jackStraw score.

To identify the neuronal cell types, each cluster was annotated with the average expression value of known marker genes for each cell type from previous atudies^57, 58^.

### Microarray

Human spinal cord tissues derived from gestational week 18 were obtained with a protocol approved by the Institutional Review Board committees at Chonnam National University Hospital and Gwangju Institute of Science and Technology. The hSCO samples were prepared based on the protocol described above. RNA purity and integrity were evaluated using an ND-1000 Spectrophotometer (NanoDrop, Wilmington, USA) and Agilent 2100 Bioanalyzer (Agilent Technologies, Palo Alto, USA). The Affymetrix Whole Transcript Expression array process was executed according to the manufacturer’s protocol (GeneChip Whole Transcript PLUS reagent Kit). cDNA was synthesized using the GeneChip WT (Whole Transcript) amplification kit as described by the manufacturer. The sense cDNA was then fragmented and biotin-labeled with TdT (terminal deoxynucleotidyl transferase) using the GeneChip WT Terminal labeling kit. Approximately 5.5 μg of labeled DNA target was hybridized to the Affymetrix GeneChip Human 2.0 ST Array at 45°C for 16 h. Hybridized arrays were washed and stained on a GeneChip Fluidics Station 450 and scanned with a GCS3000 Scanner (Affymetrix). Signal values were computed using the Affymetrix^®^ GeneChip™ Command Console software.

The data were normalized with a robust multi-average (RMA) method implemented in Affymetrix^®^ Power Tools (APT). We have focused on genes variable along with cellular differentiation toward the spinal cord development on hSCO samples. Variable genes were identified by calculating the median absolute deviation (MAD) across all hSCO samples with the cutoff of top 5% of MAD values, which resulted in 1,365 genes. For comparison with the human fetal spinal cord, Hierarchical cluster analysis was performed on the previously found 1,365 variable genes with all hSCO samples and the human fetal spinal cord sample. The variable genes were divided into two clusters, up- and down-regulated genes on differentiation, which were used for analyzing functional enrichment in the GO terms or biological processes.

### Chick notochord culture

Fertilized eggs (Pulmuone) were incubated at 37 *°C* in a humidified incubator to obtain HH 25-27. The embryos were removed from the eggs and washed in ice-cold PBS. The intact notochords were dissected and divided into several parts under a dissection microscope. Each notochord fragment was transferred into a 96-well plate containing the day 7 organoid. The organoids with chick notochord were cultured in DM containing 0.1 μM RA for 8 days. On day 15, the organoids were grown in DM supplemented with bone morphogenetic protein 4 (BMP4) (50 ng/mL, PeproTech) or Purmorphamine (1 μm, Merck) for 6 days. The medium was changed every alternate day.

### Neuromuscular junction formation

Human skeletal muscle cells (hSkMCs) were purchased from PromoCell (C-12530). The 3D myotube of the hSkMC formation protocol has been previously described^59^. Briefly, hSkMCs were seeded at a high density in the device and cultured in skeletal muscle cell growth medium (PromoCell, C-23060) for 2 days. The growth medium was then replaced with skeletal muscle cell differentiation medium (PromoCell, C-23061). After 7 days, when cells had formed a condensed myotube, the medium was switched back to the growth medium to allow for the stable differentiation of hSkMCs. On day 26, the developing hSCOs were cultured with purmorphamine (Day 15) and plated on the 3D hSkMC myotube in differentiation medium for hSCO. The co-culture medium was changed every 3 days. The hSCOs and myotubes were fixed on day 38 of the co-culture.

### Dorsal root ganglion (DRG) culture

DRGs were isolated from all levels of the spinal cord of Thy1-YFP mice (P7). After exposing the spinal cord, DRGs were dissected with a dissection microscope and collected in a petri dish filled with ice-cold HBSS. DRG explants were prepared by trimming nerve roots and removing the connective tissue sheath with microsurgical scissors. Each DRG explant was placed with a single hSCO in a round-bottom well of a 96-well plate. After 7 days, the fused hSCOs-DRG complexes were transferred to the petri dish. For all procedures, the samples were cultured in a 1:1 mixture of DMEM/F-12 and neurobasal medium as described above.

### Statistical analysis

Statistical analyses were performed using an unpaired Student’s t-test. All analyses were performed with GraphPad Prism 8 software, and the results are presented as mean ± SEM. P values <0.05 were considered statistically significant.

### Ethical statements

The human PSC study was approved by the Korea University Institutional Review Board. All animal maintenance and experimental procedures were approved by members of the Laboratory Animal Research Center at Korea University College of Medicine.

## Acknowledgments

We would like to thank the Korea Basic Science Institute, Korea Brain Research Institute, Dr. Kyung-Sook Yang and Ms. Jieun Na, in particular, for technical support. We also thank Prof. Jung Hosung (Yonsei University) for critical comments. This research was supported by the Brain Research Program through the National Research Foundation (NRF), which is funded by the Korean Ministry of Science, ICT & Future Planning (NRF-2015M3C7A1028790, NRF-2017M3C7A1047654, and NRF-2017M3A9B3061308).

## Author Contributions

**Ju-Hyun Lee:** Conceptualization, Methodology, Validation, Investigation, Visualization, Formal analysis, Writing – Original Draft, Writing – review & editing. **Hyogeun Shin:** Methodology, Investigation, Resources, Visualization. **Mohammed R. Shaker:** Validation, Investigation. **Hyun Jung Kim:** Methodology, Investigation. **June Hoan Kim:** Data Curation. **Namwon Lee:** Software, Resources, Data Curation, Formal analysis. **Minjin Kang:** Investigation, Resources. **Subin Cho:** Software, Resources, Visualization, Data Curation, Formal analysis. **Tae Hwan Kwak:** Resources. **Jong Woon Kim:** Resources. **Mi-Ryong Song:** Resources. **Seung-Hae Kwon:** Resources. **Dong Wook Han:** Resources. **Sanghyuk Lee:** Resources. **Se-Young Choi:** Conceptualization. **Im Joo Rhyu:** Resources. **Hyun Kim:** Resources. **Dongho Geum:** Conceptualization, Resources. **Il-Joo Cho:** Conceptualization, Resources. **Woong Sun:** Conceptualization, Supervision, Project administration, Funding acquisition, Writing – original draft, Writing – review & editing.

## Code availability

The code for training the deep-learning models in this study are available at https://github.com/im-namwon/stemcell-classification.

## References

1. Smith, J.L. & Schoenwolf, G.C. Neurulation: coming to closure. Trends in neurosciences 20, 510–517 (1997).

2. Jankowska, E. Spinal interneuronal systems: identification, multifunctional character and reconfigurations in mammals. The Journal of physiology 533, 31–40 (2001).

3. Lu, D.C., Niu, T. & Alaynick, W.A. Molecular and cellular development of spinal cord locomotor circuitry. Frontiers in molecular neuroscience 8, 25 (2015).

4. Juriloff, D.M. & Harris, M.J. Mouse models for neural tube closure defects. Human molecular genetics 9, 993–1000 (2000).

5. Greene, N.D., Stanier, P. & Copp, A.J. Genetics of human neural tube defects. Human molecular genetics 18, R113–R129 (2009).

6. Holmes, L.B. et al. The teratogenicity of anticonvulsant drugs. New England Journal of Medicine 344, 1132–1138 (2001).

7. Loeken, M.R. in American Journal of Medical Genetics Part C: Seminars in Medical Genetics, Vol. 135 77–87 (Wiley Online Library, 2005).

8. Matok, I. et al. Exposure to folic acid antagonists during the first trimester of pregnancy and the risk of major malformations. British journal of clinical pharmacology 68, 956–962 (2009).

9. Mølgaard-Nielsen, D. & Hviid, A. Newer-generation antiepileptic drugs and the risk of major birth defects. Jama 305, 1996–2002 (2011).

10. Warmflash, A., Sorre, B., Etoc, F., Siggia, E.D. & Brivanlou, A.H. A method to recapitulate early embryonic spatial patterning in human embryonic stem cells. Nature methods 11, 847–854 (2014).

11. Haremaki, T. et al. Self-organizing neuruloids model developmental aspects of Huntington’s disease in the ectodermal compartment. Nature biotechnology 37, 1198–1208 (2019).

12. Moris, N. et al. An in vitro model of early anteroposterior organization during human development. Nature, 1–6 (2020).

13. Kim, J., Koo, B.-K. & Knoblich, J.A. Human organoids: model systems for human biology and medicine. Nature Reviews Molecular Cell Biology 21, 571–584 (2020).

14. Lancaster, M.A. et al. Cerebral organoids model human brain development and microcephaly. Nature 501, 373 (2013).

15. Jo, J. et al. Midbrain-like organoids from human pluripotent stem cells contain functional dopaminergic and neuromelanin-producing neurons. Cell stem cell 19, 248–257 (2016).

16. Qian, X. et al. Brain-region-specific organoids using mini-bioreactors for modeling ZIKV exposure. Cell 165, 1238–1254 (2016).

17. Quadrato, G. et al. Cell diversity and network dynamics in photosensitive human brain organoids. Nature 545, 48 (2017).

18. Paşca, A.M. et al. Functional cortical neurons and astrocytes from human pluripotent stem cells in 3D culture. Nature methods 12, 671 (2015).

19. Kadoshima, T. et al. Self-organization of axial polarity, inside-out layer pattern, and speciesspecific progenitor dynamics in human ES cell–derived neocortex. Proceedings of the National Academy of Sciences 110, 20284–20289 (2013).

20. Meinhardt, A. et al. 3D reconstitution of the patterned neural tube from embryonic stem cells. Stem cell reports 3, 987–999 (2014).

21. Ogura, T., Sakaguchi, H., Miyamoto, S. & Takahashi, J. Three-dimensional induction of dorsal, intermediate and ventral spinal cord tissues from human pluripotent stem cells. Development 145, dev162214 (2018).

22. Hor, J.H. et al. Cell cycle inhibitors protect motor neurons in an organoid model of Spinal Muscular Atrophy. Cell death & disease 9, 1–12 (2018).

23. Martins, J.-M.F. et al. Self-organizing 3D human trunk neuromuscular organoids. Cell stem cell 26, 172–186. e176 (2020).

24. Rifes, P. et al. Modeling neural tube development by differentiation of human embryonic stem cells in a microfluidic WNT gradient. Nature Biotechnology, 1–9 (2020).

25. Quadrato, G., Brown, J. & Arlotta, P. The promises and challenges of human brain organoids as models of neuropsychiatric disease. Nature medicine 22, 1220 (2016).

26. Velasco, S. et al. Individual brain organoids reproducibly form cell diversity of the human cerebral cortex. Nature, 1 (2019).

27. Denham, M. et al. Multipotent caudal neural progenitors derived from human pluripotent stem cells that give rise to lineages of the central and peripheral nervous system. Stem Cells 33, 1759–1770 (2015).

28. Anderson, M.J. et al. TCreERT2, a transgenic mouse line for temporal control of Cre-mediated recombination in lineages emerging from the primitive streak or tail bud. PloS one 8, e62479 (2013).

29. Nagai, T. et al. The Expression of the MouseZic1, Zic2, andZic3Gene Suggests an Essential Role forZicGenes in Body Pattern Formation. Developmental biology 182, 299–313 (1997).

30. Aaku-Saraste, E., Hellwig, A. & Huttner, W.B. Loss of occludin and functional tight junctions, but not ZO-1, during neural tube closure—remodeling of the neuroepithelium prior to neurogenesis. Developmental biology 180, 664–679 (1996).

31. Nishimura, T., Honda, H. & Takeichi, M. Planar cell polarity links axes of spatial dynamics in neural-tube closure. Cell 149, 1084–1097 (2012).

32. Bretzner, F. & Brownstone, R.M. Lhx3-Chx10 reticulospinal neurons in locomotor circuits. Journal of Neuroscience 33, 14681–14692 (2013).

33. Floyd, T.L., Dai, Y. & Ladle, D.R. Characterization of calbindin D28k expressing interneurons in the ventral horn of the mouse spinal cord. Developmental Dynamics 247, 185–193 (2018).

34. Dale, N. Reciprocal inhibitory interneurones in the Xenopus embryo spinal cord. The Journal of Physiology 363, 61–70 (1985).

35. Shin, H. et al. Multifunctional multi-shank neural probe for investigating and modulating long-range neural circuits in vivo. Nature communications 10, 1–11 (2019).

36. Suzue, T. Respiratory rhythm generation in the in vitro brain stem-spinal cord preparation of the neonatal rat. The Journal of physiology 354, 173–183 (1984).

37. Hanson, M.G. & Landmesser, L.T. Characterization of the circuits that generate spontaneous episodes of activity in the early embryonic mouse spinal cord. Journal of Neuroscience 23, 587–600 (2003).

38. Saito, A. et al. Modulation of neuronal network activity using magnetic nanoparticle-based astrocytic network integration. Biomater Sci 3, 1228–1235 (2015).

39. Zafeiriou, M.-P et al. Developmental GABA polarity switch and neuronal plasticity in Bioengineered Neuronal Organoids. Nature Communications 11, 1–12 (2020).

40. Greene, N.D. & Copp, A.J. Neural tube defects. Annual review of neuroscience 37, 221–242 (2014).

41. Agopian, A., Tinker, S.C., Lupo, P.J., Canfield, M.A. & Mitchell, L.E. Proportion of neural tube defects attributable to known risk factors. Birth Defects Research Part A: Clinical and Molecular Teratology 97, 42–46 (2013).

42. Weston, J. et al. Monotherapy treatment of epilepsy in pregnancy: congenital malformation outcomes in the child. Cochrane Database of Systematic Reviews (2016).

43. Kawada, J. et al. Generation of a motor nerve organoid with human stem cell-derived neurons. Stem cell reports 9, 1441–1449 (2017).

44. Sternfeld, M.J. et al. Speed and segmentation control mechanisms characterized in rhythmically-active circuits created from spinal neurons produced from genetically-tagged embryonic stem cells. Eiife 6, e21540 (2017).

45. Zheng, Y. et al. Dorsal-ventral patterned neural cyst from human pluripotent stem cells in a neurogenic niche. Science advances 5, eaax5933 (2019).

46. Davidson, B.P., Kinder, S.J., Steiner, K., Schoenwolf, G.C. & Tam, P.P. Impact of node ablation on the morphogenesis of the body axis and the lateral asymmetry of the mouse embryo during early organogenesis. Developmental biology 211, 11–26 (1999).

47. Stemple, D.L. Structure and function of the notochord: an essential organ for chordate development. Development 132, 2503–2512 (2005).

48. Moury, J.D. & Schoenwolf, G.C. Cooperative model of epithelial shaping and bending during avian neurulation: autonomous movements of the neural plate, autonomous movements of the epidermis, and interactions in the neural plate/epidermis transition zone. Developmental dynamics 204, 323–337 (1995).

49. Sato, T. et al. Single Lgr5 stem cells build crypt–villus structures in vitro without a mesenchymal niche. Nature 459, 262 (2009).

50. Yoon, S.-J. et al. Reliability of human cortical organoid generation. Nature methods 16, 75–78 (2019).

51. Shin, H. et al. Neural probes with multi-drug delivery capability. Lab on a Chip 15, 3730–3737 (2015).

52. Lee, Y.J., Song, K.-I., Kang, J.Y. & Lee, S.H. in 2015 37th Annual International Conference of the IEEE Engineering in Medicine and Biology Society (EMBC) 3415–3418 (IEEE, 2015).

53. Bakkum, D.J. et al. Parameters for burst detection. Frontiers in computational neuroscience 7, 193 (2014).

54. Dobin, A. et al. STAR: ultrafast universal RNA-seq aligner. Bioinformatics 29, 15–21 (2013).

55. Butler, A., Hoffman, P., Smibert, P., Papalexi, E. & Satija, R. Integrating single-cell transcriptomic data across different conditions, technologies, and species. Nature biotechnology 36, 411–420 (2018).

56. Hafemeister, C. & Satija, R. Normalization and variance stabilization of single-cell RNA-seq data using regularized negative binomial regression. Genome biology 20, 1–15 (2019).

57. Rosenberg, A.B. et al. Single-cell profiling of the developing mouse brain and spinal cord with split-pool barcoding. Science 360, 176–182 (2018).

58. Delile, J. et al. Single cell transcriptomics reveals spatial and temporal dynamics of gene expression in the developing mouse spinal cord. Development 146 (2019).

59. Sakar, M.S. et al. Formation and optogenetic control of engineered 3D skeletal muscle bioactuators. Lab on a Chip 12, 4976–4985 (2012).

