## Supplemental information for "Human spinal cord organoids exhibiting neural tube morphogenesis for a quantifiable drug screening system of neural tube defects"

### Supplementary Figures

Supplementary Figure 1_Related to Figure 1.

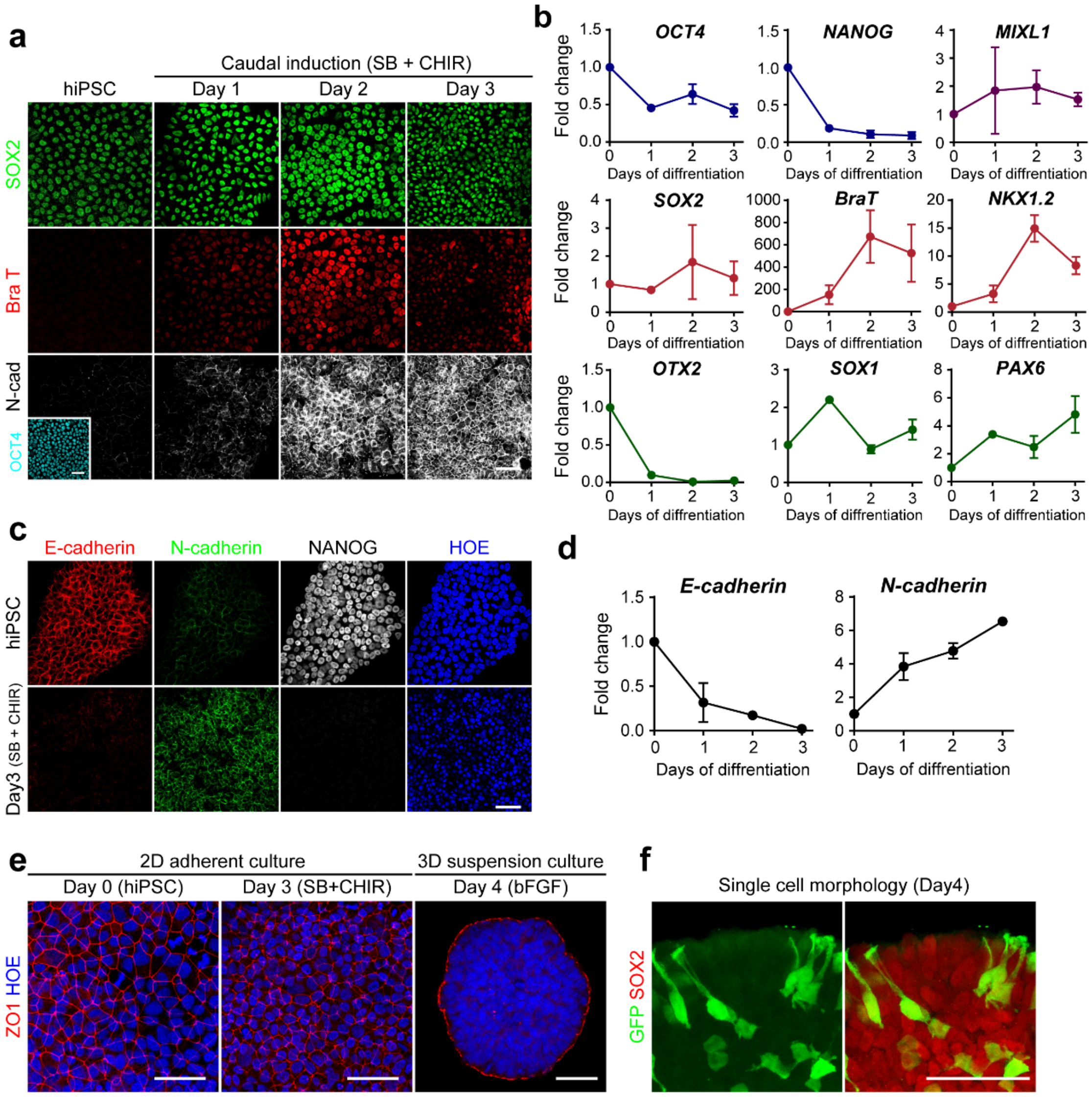


**Supplementary Fig. 1 (related to Fig. 1). Generation of caudal NSCs from hiPSCs in 2D**

**a.** Caudal induction (SB + CHIR) of hiPSCs on 2D culture showed transient induction of the NMp marker BraT and subsequent downregulation of BraT, leading to the conversion of hiPSCs into N-cad positive neural stem cells. **b**. Real-time PCR profiles of gene expression for pluripotency marker (OCT4 and NANOG), mesendoderm (MIXL1), neuromesoderm (SOX2, BraT, and NKX1.2), neuroectoderm (SOX1 and PAX6), and rostral neuroectoderm (OTX2). **c**. The E- to N-cadherin switch was present during 2D induction. Immunofluorescence analysis of E-cadherin (red), N-cadherin (green), and Nanog (white). Nuclei were counterstained with Hoechst (blue). **d.** Relative expression of the E- and N- cadherin mRNAs depending on the days of differentiation. **e.** The maintenance of apical polarity from 2D to 3D culture. During the 3D conversion process, surface neuroepithelia cells (NE) re-established their polarity within 24 hours. Apical polarity was visualized with ZO1 immunolabeling (red). Nuclei were counterstained with Hoechst (blue). **f.** The pseudostratified morphology of NE cells on day 4 (bFGF, day1). Note that the internal neural stem cells (SOX2+) exhibit distinct cell morphology. Cell morphologies was visualized with a small fraction of GFP-labeled cells initially mixed with non-labeled hPSCs. All scale bars, 50 μm.

Supplementary Figure 2_Related to Figure 1.

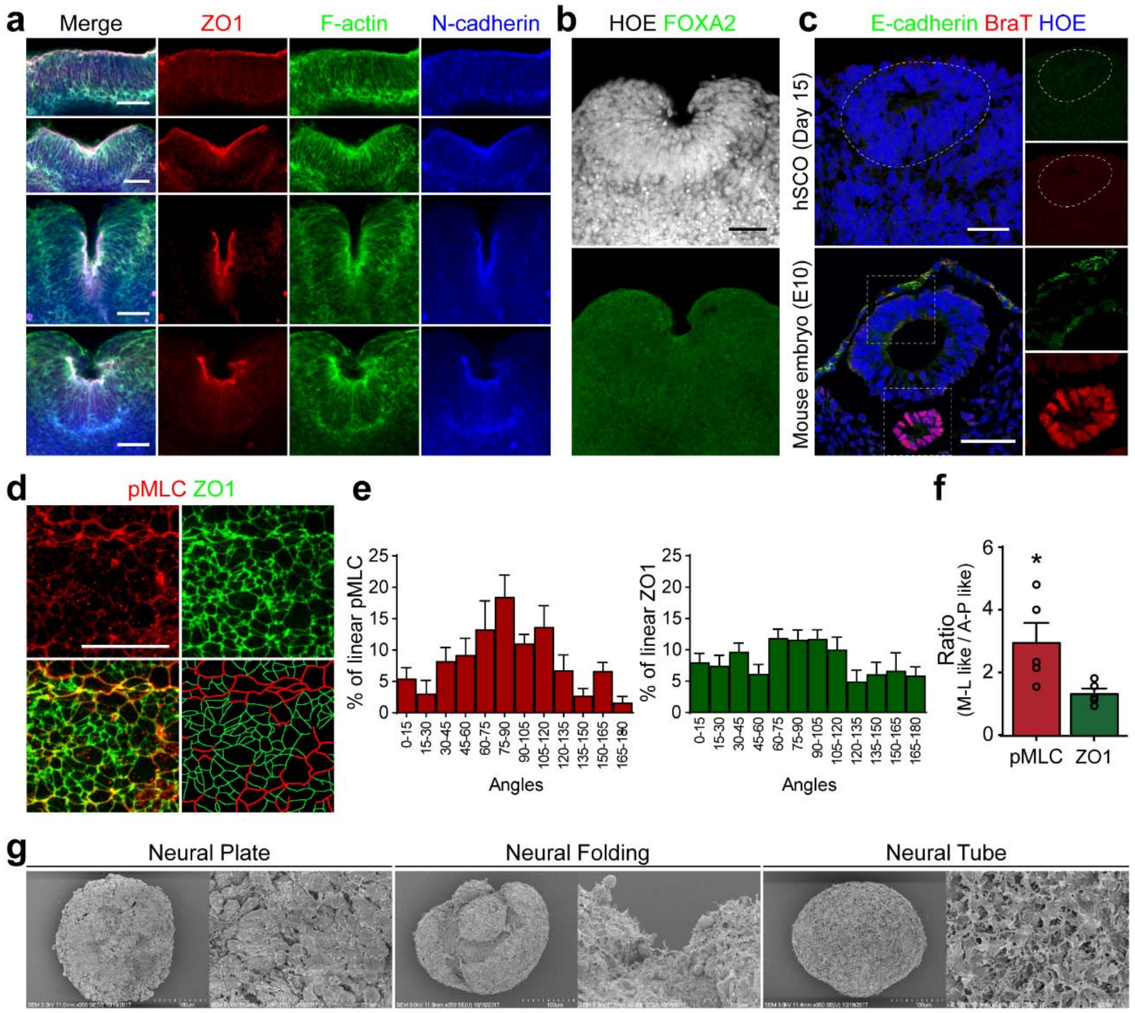


**Supplementary Fig. 2 (related to Fig. 1). hSCOs mimicking neurulation morphogenesis *in vivo***

**a.** Morphology of neural folding at different stages of development. **b.** Immunofluorescence staining of the floor plate marker FOXA2 (green) showed the absence of floor plate cells at the neural groove. **c.** Absence of non-neural tissue components in the hSCOs. The non-neural epithelium was visualized with E-cadherin (green) and the mesodermal tissue was visualized with BraT (red) staining. Developing mouse neural tube (bottom) is shown as a control. **d.** High-magnification image of the planar cell polarity (PCP) region. The right-bottom image shows traces of pMLC and ZO1 expression. **e.** Quantification of PCP with angular analysis of pMLC linearity (left) and ZO1 linearity (right). To measure the angles of pMLC and ZO1 cables, we selected a line linking two or more cells. Compared to ZO1 cables, the pMLC cables tend to align laterally. **f.** The ratio of M-L like axis (from 45 to 135 degrees) to A-P like axis (from 0 to 45 and from 135 to 180 degrees). (Unpaired *t*-test; **P 0.0326, n=5) **g.** SEM images of hSCOs at different stages of development. Note that the surfaces of the first spheroids were relatively smooth with strong tight junctions, while NT-stage hSCOs had rough surfaces. All scale bars, 50 μm.

Supplementary Figure 3_Related to Figure 1.

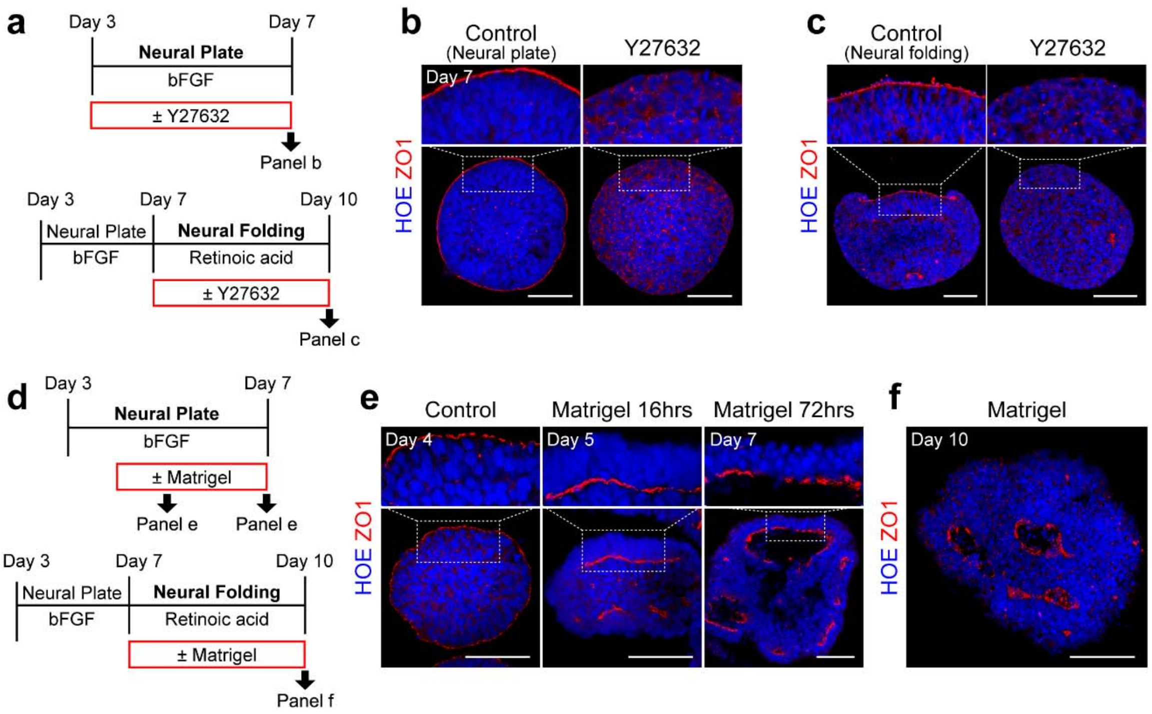


**Supplementary Fig 3 (related to Fig. 1). Effects of perturbation of apico-basal polarity on neural tube morphogenesis.**

**a.** Treatment schemes of Y-27632 (Rock inhibitor; 10 μm) for panels (b, top) and (c, bottom). **b.** The effects of Y-27632 on the establishment of neural plate. The apical polarity at neural plate-stage was visualized with ZO1 (red). Note that the Y-27632 treatments disrupted apical localization of ZO1. **c.** The effects of Y-27632 at the neural folding-stage. The apical polarity at neural plate-stage was visualized with ZO1 (red). Note that the neural folding was not induced by the Y-27632 treatments and the organoids exhibited round morphology. **d.** Treatment scheme for Matrigel-embedding experiments on panels (e, top) and (f, bottom). **e.** Matrigel embedding at early the neural plate-stage rapidly reversed the apical polarity, and promoted cavitation (by 72 hr). The apical polarity at neural plate-stages was visualized with ZO1 (red). **f.** Matrigel embedding at the neural folding-stage also resulted in the ventricle-like morphogenesis as previously reported with forebrain organoids[1]. The apical polarity at neural plate-stages was visualized with ZO1 (red). The nuclei were counterstained with Hoechst (blue) in all panels. All scale bars, 50 μm.

Supplementary Figure 4_Related to Figure 1&3.

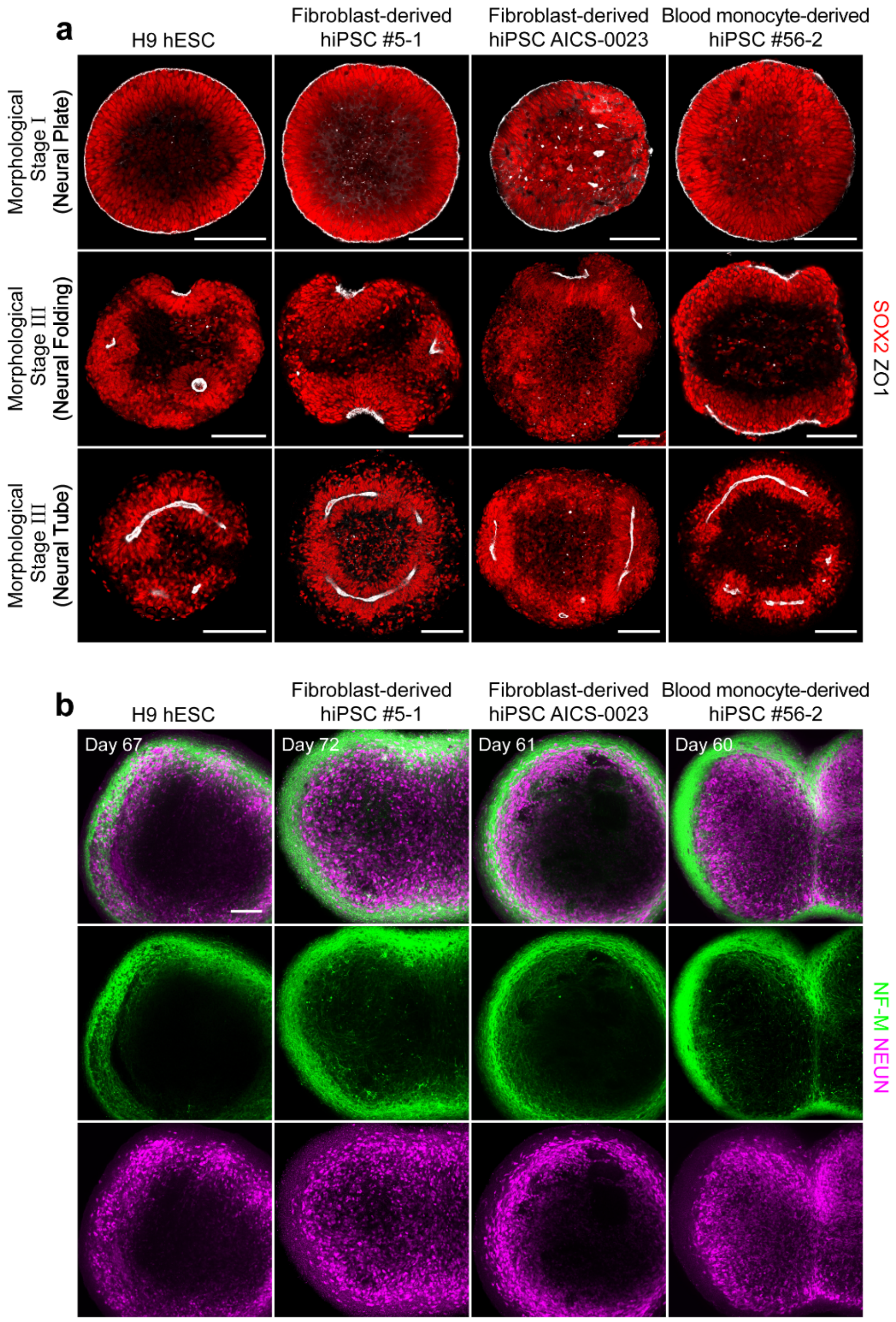


**Supplementary Fig. 4 (related to Fig. 1, 3). Morphogenesis of hSCOs from various hPSC lines**

**a-b.** Early tube morphogenesis of hSCOs from H9 hESCs (left), two different epidermal fibroblast-derived hiPSC lines (middle), and peripheral blood monocyte-derived hiPSCs (right). Each line shows three different stages of neural tube morphogenesis, visualized with staining for SOX2 (red) and ZO1 (white) **(a).** Mature morphology of hSCOs from different hPSC lines consistently exhibited core-to-shell organization of neuronal cell bodies and neurites (**b).**

Supplementary Figure 5_Related to Figure 2.

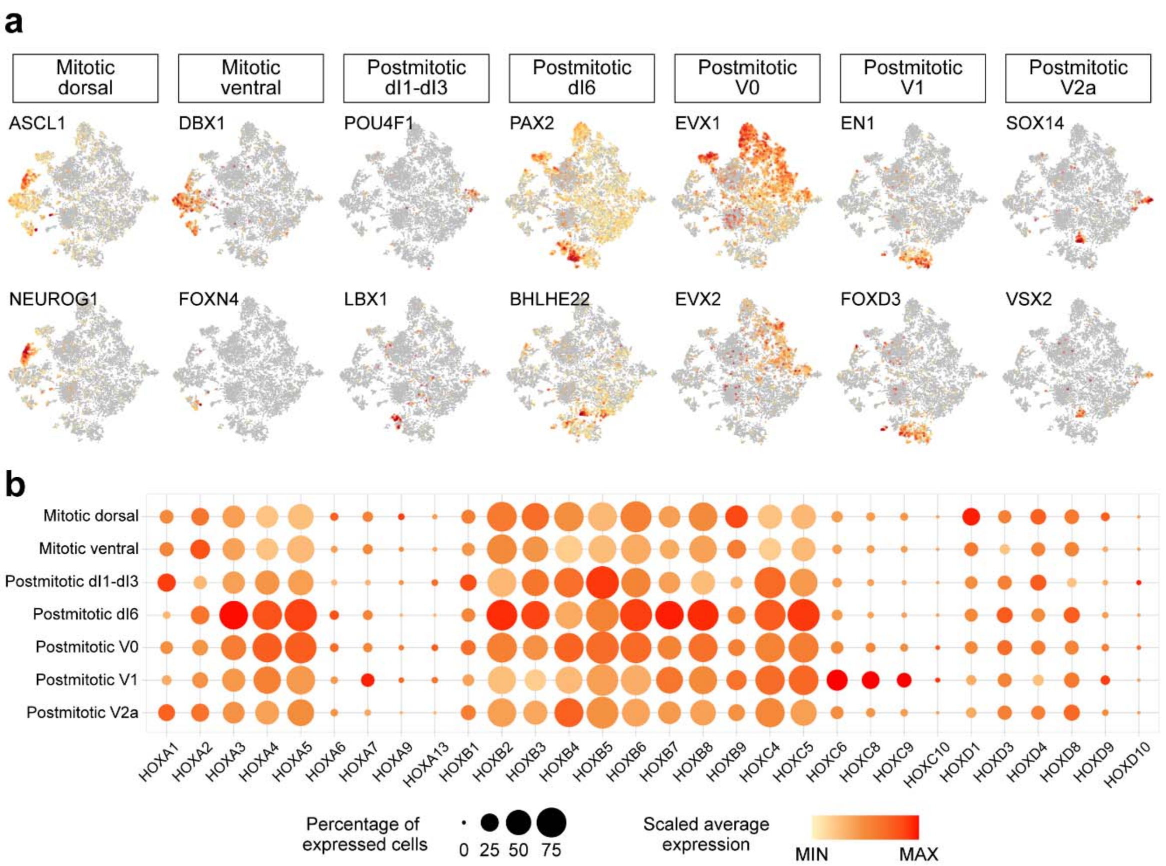


**Supplementary Fig. 5 (related to Fig. 2). Analysis of single-cell RNA sequencing for dorsoventral and anterior-posterior specifications in 1-month hSCOs**

**a.** t-SNE plots showing the clustering of major dorsoventral region-specific gene expression levels. **b.** Dot plots showing the expression of HOX genes across the 7 sub-clusters present in Figure 2c. All sub-clusters exhibit a similar pattern of HOX genes expression similar to the cervical-thoracic identity. Circle size and color represent the percentage of expressed cells and the average of gene expression, respectively.

Supplementary Figure 6_Related to Figure 2.

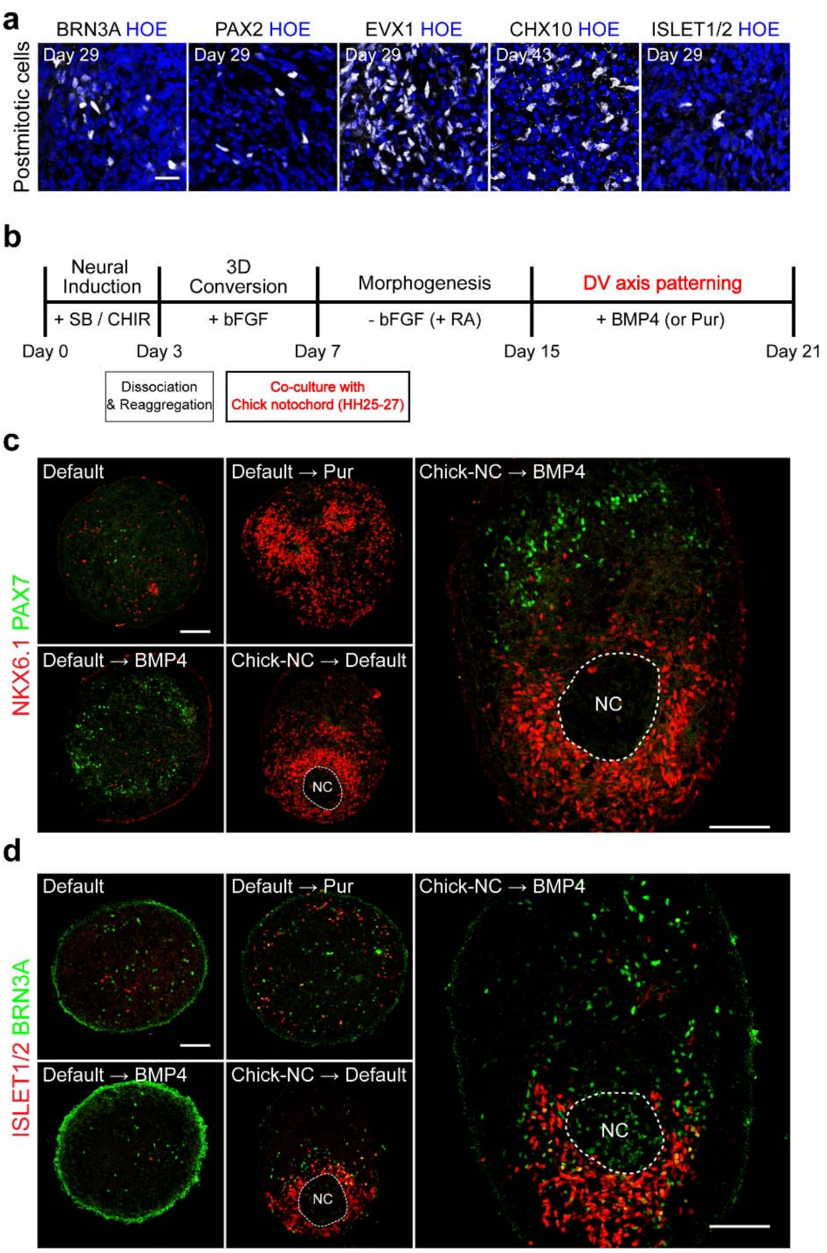


**Supplementary Fig. 6 (related to Fig. 2). Dorsoventral patterning in hSCOs**

**a.** Spinal interneurons in hSCOs. Markers were used for dI 1-3 neurons (POU4F1), dI 4-6 / V0-1 neurons (PAX2), V0 neurons (EVX1), V2 neurons (VSX2), and motor neurons (ISLET1/2).

Scale bars, 20 μm. Note that these domain-specific markers were rather scattered in the hSCOs. **b.** Induction of dorsoventral axis by co-culture with chick notochord. Schematics for the co-culture of hSCOs and chick embryonic notochord. **c-d.** Dorsoventral axis formation within hSCOs. Control hSCOs did not exhibit overt dorsoventral patterning as revealed either by NKX6.1 (red) / PAX7 (green) double staining (c) or ISLET1/2 (red) / BRN3A (green) double staining (d). When hSCOs were treated with SHH agonist Purmophamine (Pur) or BMP4 on day 15, hSCOs appeared to obtain ventral or dorsal identities, respective. When chick notochord (Chick-NC) was co-cultured with hSCOs from day 7 (neural plate-stage), notochord acted as ventralizing center and the adjacent to the notochord area of hSCOs exhibited ventral marker expressions. Finally, the combination of notochord co-culture and BMP4 treatments (Chick-NC + BMP4) resulted in the dorsoventral patterning within the hSCOs. NC, notochord. Scale bars, 100 μm

Supplementary Figure 7_Related to Figure 2.

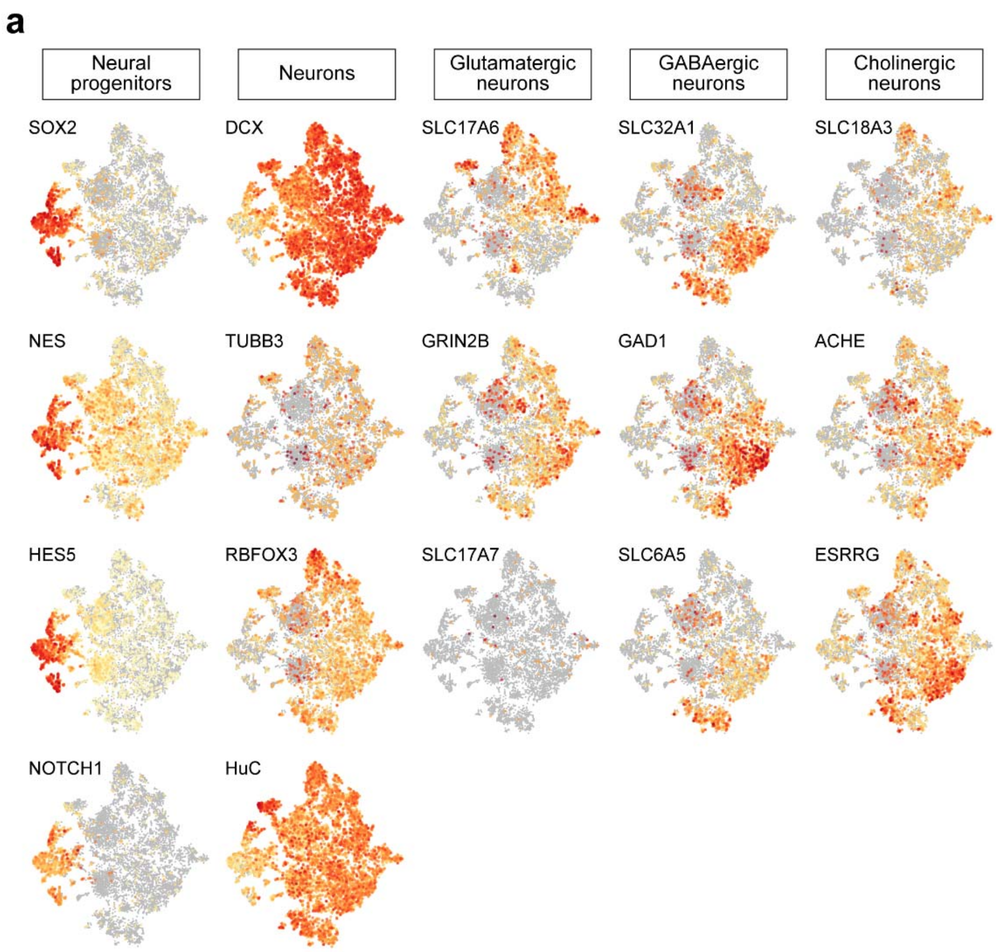


**Supplementary Fig. 7 (related to Fig. 2). Analysis of single-cell RNA sequencing for neuronal cell type specification in 1-month hSCOs**

- **a.** t-SNE plots showing cell-type-specific gene expression levels; neural progenitors, differentiated neurons, and neuronal subtype defined by neurotransmitters.

Supplementary Figure 8_Related to Figure 3.

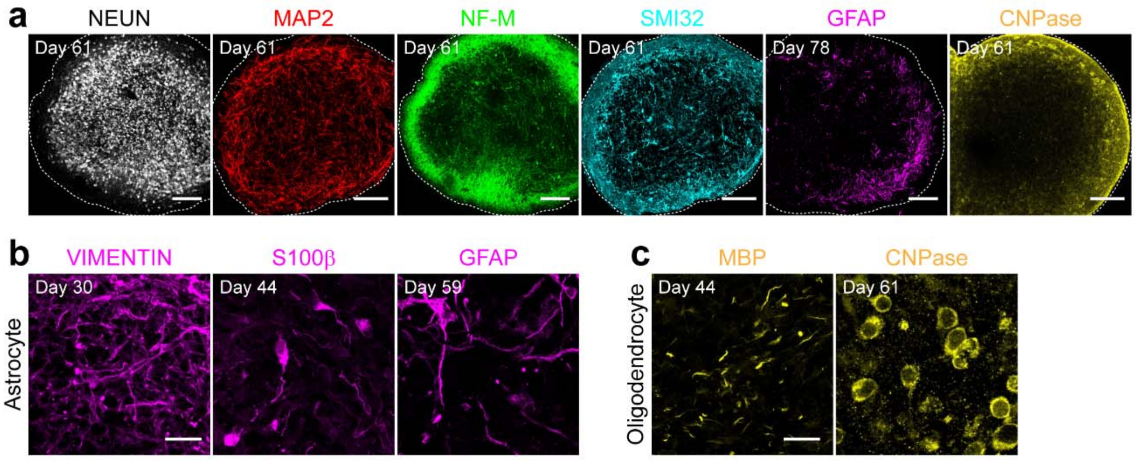


**Supplementary Fig. 8 (related to Fig. 3). Immunostaining of various cell types in hSCOs with prolonged culture**

**a**. The core-to-shell organization of hSCOs. Markers were used for neuronal nuclei (NEUN), dendrites (MAP2), axon (NF-M and SMI32), astrocytes (GFAP), and oligodendrocytes (CNPase). Scale bar, 200 μm. **b-c.** Gliogenesis in hSCOs. Markers were used for immature (vimentin), intermediate (S100β), and mature (GFAP) astrocytes (**b**); mature oligodendrocytes, MBP, and CNPase (**c**). Scale bar, 20 μm.

Supplementary Figure 9_Related to Figure 3.

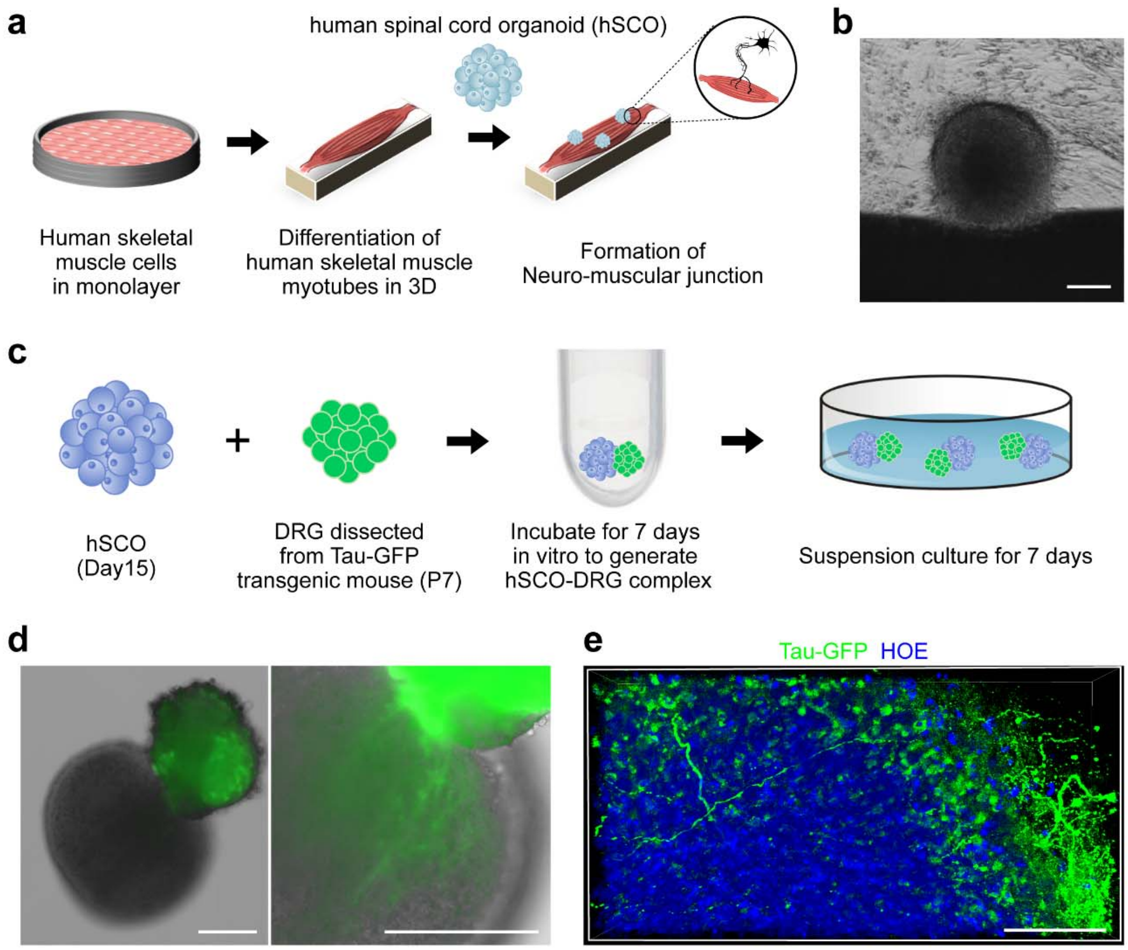


**Supplementary Fig. 9 (related to Fig. 3). Co-culture of hSCOs with peripheral counterparts**

**a**. Schematics for the co-culture of hSCOs and a human myotube. **b**. Brightfield image of hSCOs attaching to the myotube. Scale bar, 200 μm. **c.** Schematics for the co-culture of hSCOs and a mouse DRG. The DRG was obtained from P7 Tau-GFP transgenic mice and co-cultured with day-15 hSCOs in 96-well plates with U-shaped bottom for 7 days. After visual confirmation of the strong fusion of the hSCOs and DRG, the fused hSCOs were transferred to an uncoated culture dish and subjected to another 7 days of suspension culture. **d.** Brightfield image of fused hSCOs with DRG (green). The right image shows the robust entry of GFP-labeled fibers into hSCOs. Scale bar, 200 μm. **e.** High-resolution image of GFP-labeled sensory fiber into the hSCO territory. Scale bar, 50 μm.

Supplementary Figure 10_Related to Figure 4.

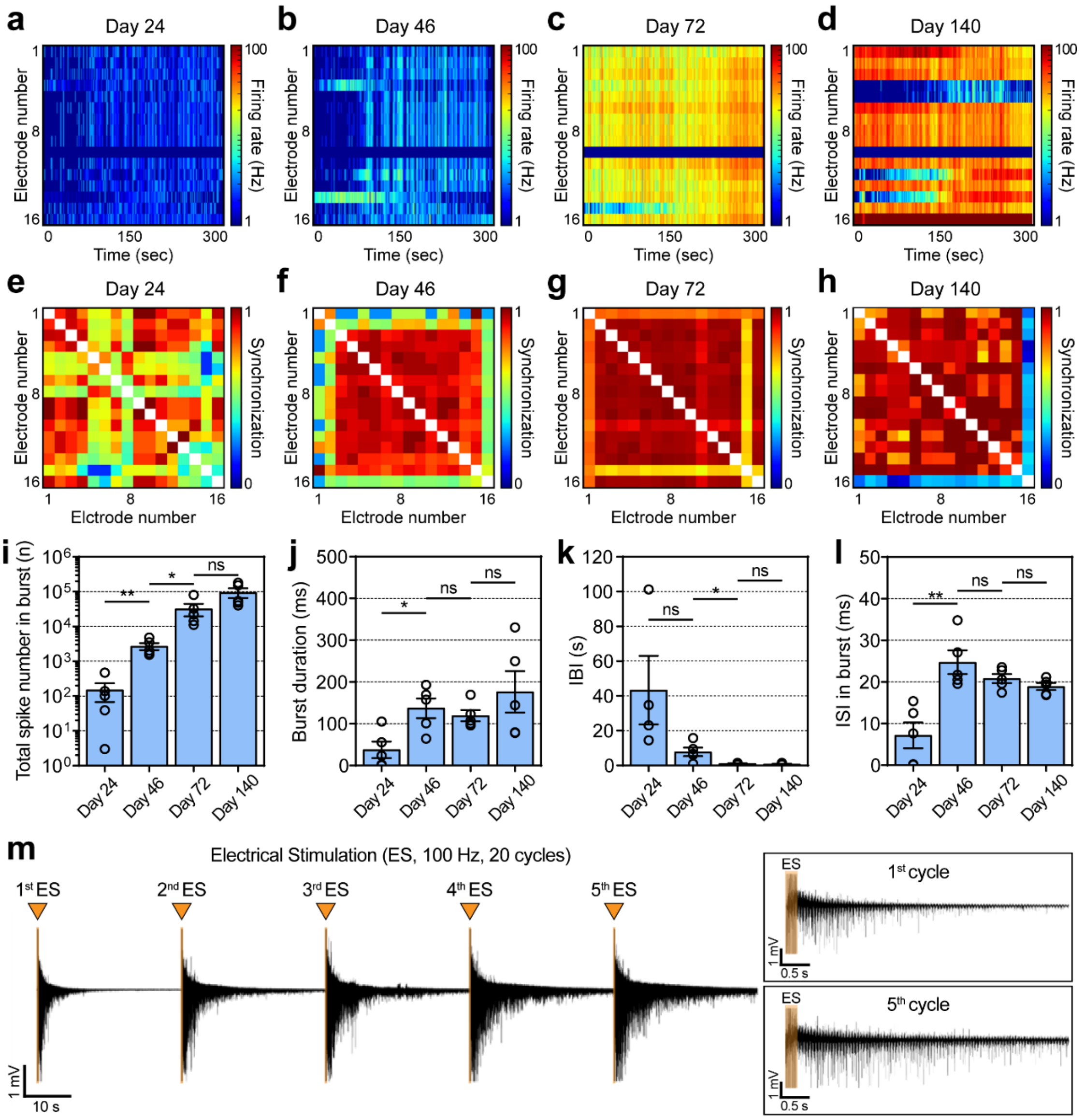


**Supplementary Fig. 10 (related to Fig. 4). Changes in the patterns of neural activities in hSCOs**

**a-d.** Representative heatmaps showing neural activities at Day 24 **(a)**, Day 46 **(b)**, Day 72 **(c)**, and Day 140 **(d)**. **e-h.** Representative cross-correlation matrices showing synchronization between signal-recorded electrodes at Day 24 **(e)**, Day 46 **(f)**, Day 72 **(g)**, and Day 140 **(h)**. **i-j.** Changes in the patterns of burst activity as the hSCOs mature (error bars indicate s.e.m. n=5). Total spike number in burst activities (Day 24 – 46: ** P 0.0035; Day 46 – 72: * P 0.0483; Day 72 – 140: ns P 0.0858; n=number of the samples, Two-tailed unpaired *t*-test) **(i).** Burst duration (Day 24 – 46: * P 0.0119; Day 46 – 72: ns P 0.5309; Day 72 – 140: ns P 0.2977; n=number of the samples, Two-tailed unpaired *t*-test) **(j).** Inter burst interval (IBI) (Day 24 – 46: ns P 0.0824; Day 46 – 72: * P 0.0246; Day 72 – 140: ns P 0.2702; n the number of the samples, Two-tailed unpaired *t*-test) **(k).** Inter spike interval (ISI) in burst activities (Day 24 – 46: ** P 0.0031; Day 46 – 72: ns P 0.2322; Day 72 – 140: ns P 0.2074; n=number of the samples, Two-tailed unpaired *t*-test) **(l). m.** Representative transient plot showing electrically evoked activities in matured hSCOs and expanded plots of the first and fifth stimulus.

Supplementary Figure 11_Related to Figure 5.

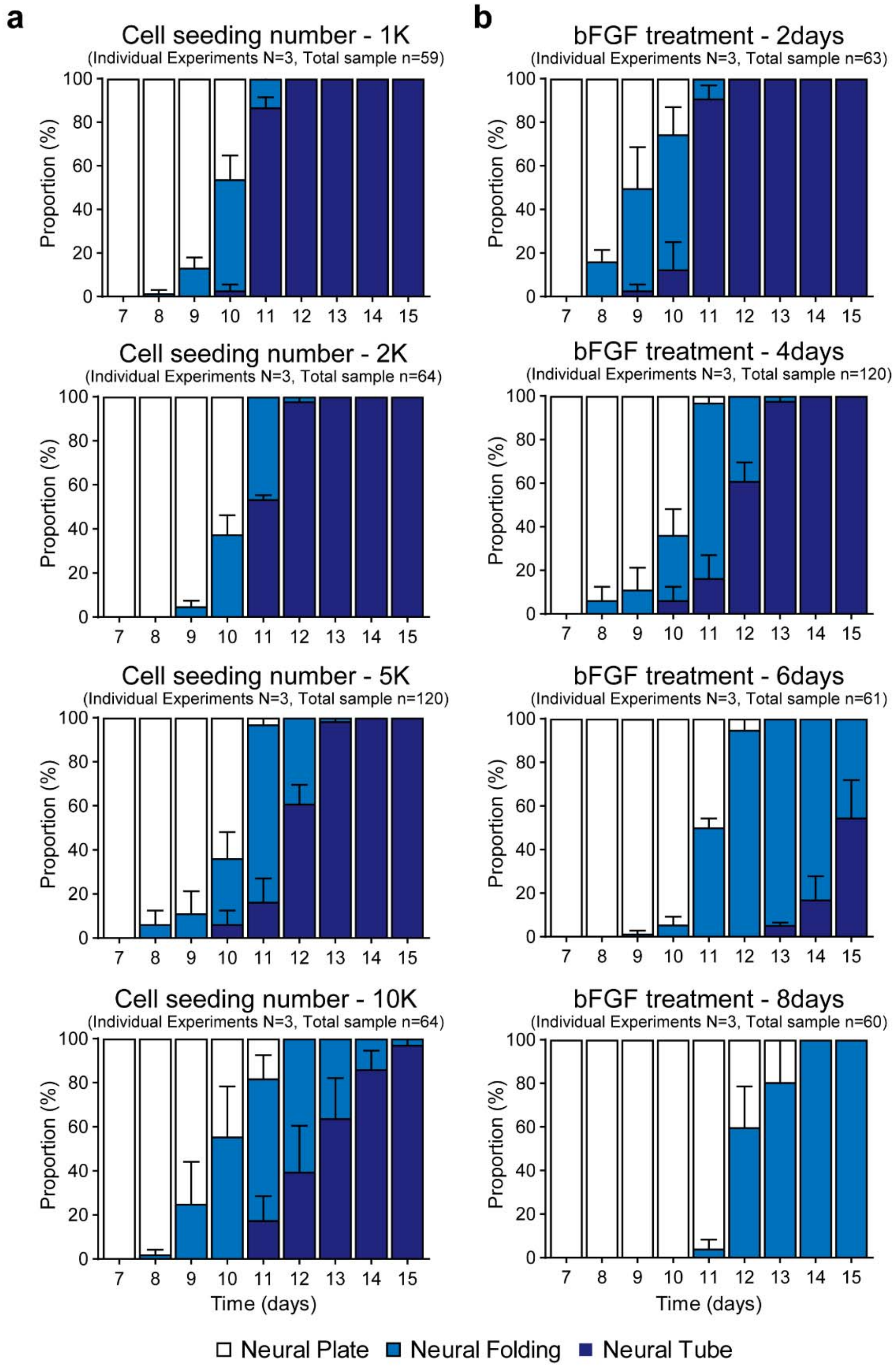

**Supplementary Fig. 11 (related to Fig. 5). Size and bFGF duration-dependent morphogenesis in hSCOs**

**a.** Quantification of cell seeding number-dependent morphogenesis. The proportion of hSCOs at indicated morphological stages (white, neural plate stage; blue, neural folding stage; navy, neural tube stage) at the indicated culture time. The hSCOs with increasing numbers of initial cell seeding spent longer time to complete tube morphogenesis. **b.** Quantification of bFGF treatment period-dependent morphogenesis. Data are expressed as mean ± S.E.M of three independent experiments. Total number of organoids analyzed in each group is shown on the top of each graph.

#### Supplementary Tables

**Supplementary Table S1. Antibodies used for immunohistochemistry**

| Antigen | Host Species | Source | Cat# | Dilution (3D) | Dilution (2D) |
|---|---|---|---|---|---|
| SOX2 | Rabbit | Millipore | AB5603 | 1:300 | 1:500 |
| SOX2 | Mouse | Santa Cruz | sc-365823 | 2.5:300 | 1:200 |
| SOX2 | Goat | Santa Cruz | sc-17320 | 2:300 | 1:250 |
| ZO1 | Rabbit | Invitrogen | 61-7300 | 2.5:300 | 1:200 |
| ZO1 | Mouse | Invitrogen | 33-9100 | 2.5:300 | 1:200 |
| GFP | Chick | Abcam | ab13970 | 1:600 | 1:1000 |
| ZIC2 | Rabbit | Millipore | AB15392 | 2.5:300 | 1:200 |
| pMLC | Rabbit | Cell signaling | 3671 | 2.5:300 | 1:200 |
| Laminin | Rabbit | Sigma | L9393 | 1:600 | 1:1000 |
| N-cadherin | Mouse | BD Biosciences | 610921 | 1:300 | 1:500 |
| a-tubulin | Mouse | Santa Cruz | sc-8035 | 1:300 | 1:500 |
| Occludin | Rabbit | Invitrogen | 71-1500 | 2.5:300 | 1:200 |
| Occludin | Mouse | Invitrogen | 33-1500 | 2.5:300 | 1:200 |
| Phospho-histone H3 | Rabbit | Upstate | 06-570 | 1:600 | 1:1000 |
| NeuN | Rabbit | Millipore | ABN78 | 1:600 | 1:1000 |
| NeuN | Mouse | Millipore | MAB377 | 1:300 | 1:500 |
| T | Goat | R&D | AF2085 | 1:300 | 1:500 |
| OCT4 | Mouse | Santa Cruz | sc-5279 | 1:300 | 1:500 |
| E-cadherin | Rabbit | Cell signaling | 3195 | 2.5:300 | 1:200 |
| NANOG | Goat | R&D | AF1977 | 2.5:300 | 1:200 |
| FOXA2 | Mouse | DSHB | 4C7 | 5:300 | 1:100 |
| Neurofilament-M | Mouse | DSHB | 2H3 | 2:300 | 1:250 |
| GFAP | Rat | Invitrogen | 13-0300 | 1:300 | 1:500 |
| MBP | Rabbit | Abcam | ab40390 | 1:600 | 1:1000 |
| LHX3 | Mouse | DSHB | 67.4E12 | 5:300 | 1:100 |
| CHX10 | Sheep | Millipore | AB9016 | 2.5:300 | 1:200 |
| GABA | Rabbit | Sigma | A2502 | 1:600 | 1:1000 |
| vGLUT1 | Guinea pig | Millipore | Ab5905 | 2:300 | 1:250 |
| Calbindin | Mouse | Swant | 300 | 1:600 | 1:1000 |
| Calretinin | Goat | Swant | CG1 | 1:300 | 1:500 |
| Parvalbumin | Rabbit | Swant | PV27 | 1:600 | 1:1000 |
| ChaT | Goat | Milipore | AB144p | 5:300 | 1:100 |

| | | | | | |
|---|---|---|---|---|---|
| VAChT | Rabbit | Synaptic Systems | 139 103 | 1:300 | 1:500 |
| Synapsin | Rabbit | Abcam | ab8 | 1:600 | 1:1000 |
| β-Tubulin III (Tuj1) | Mouse | Sigma | T8660 | 1:600 | 1:1000 |
| PSD95 | Mouse | Thermo | MA1-045 | 2:300 | 1:250 |
| MAP2 | Chick | Millipore | AB5543 | 1:3000 | 1:5000 |
| SMI32 | Mouse | Covance | SMI-32P | 1:600 | 1:1000 |
| CNPase | Mouse | Sigma | C5922 | 1:300 | 1:500 |
| Vimentin | Goat | Millipore | AB1620 | 1:300 | 1:500 |
| S100β | Mouse | Sigma | S2532 | 1:300 | 1:500 |
| PAX7 | Mouse | DSHB | PAX7-s | 2.5:30 | 1:20 |
| IRX3 | Goat | Santa Cruz | sc-22581 | 1:30 | 1:50 |
| NKX6.1 | Goat | Santa Cruz | sc-15027 | 2.5:300 | 1:200 |
| OLIG2 | Rabbit | IBL | JP18953 | 1:300 | 1:500 |
| BRN3A | Goat | Santa Cruz | sc-31984 | 1:300 | 1:500 |
| PAX2 | Mouse | DSHB | PCRP-PAX2-1A7 | 2:30 | 1:25 |
| EVX1 | Mouse | DSHB | 99.1-3A2 | 2:30 | 1:25 |
| ISLET1.2 | Mouse | DSHB | 39.4D5 | 2.5:300 | 1:200 |
| Alexa Fluor 488 Phalloidin | - | Thermo | A12379 | 1:300 | 1:500 |
| Alexa Fluor 647 Phalloidin | - | Thermo | A22287 | 1:30 | 1:50 |
| Alexa Fluor 594 conjugate a-Bungarotoxin | - | Invitrogen | B13423 | 1:300 | 1:500 |

**Supplementary Table S2. Primer sequences used for RT-PCR**

| Gene | Primer (Forward) | Primer (Reverse) | Size |
|---|---|---|---|
| OCT4 | CTGGTTCGCTTTCTCTTTCG | CTTTGAGGCTCTGCAGCTTA | 150 |
| NANOG | AAGGCCTCAGCACCTACCTA | TGCACCAGGTCTGAGTGTTC | 181 |
| MIXL1 | TTTTCTCCCCTCTTCCAGGTA | TGGGGCTTCAGACATTTCGT | 146 |
| SOX2 | GGAAAGTTGGGATCGAACAA | GCGAACCATCTCTGTGGTCT | 145 |
| Bra T | GGGTACTCCCAATCCTATTCTGAC | ACTGACTGGAGCTGGTAGGT | 148 |
| NKX1.2 | GACCCACAGAAATTCACCCG | ATCAGGGGTCTCCAGCTGTC | 138 |
| SOX1 | ACTTTTATTTCTCGGCCCGT | GGAATGGGAGGACAGGATTT | 113 |
| PAX6 | TCCGTTGGAACTGATGGAGT | TAAGGATGTTGAACGGGCAG | 146 |
| OTX2 | CCAGACATCTTCATGCGAGAG | GGCAGGTCTCACTTTGTTTTG | 147 |
| CDH1 (E-cad) | CGAGAGCTACACGTTCACGG | GGGTGTCGAGGGAAAAATAGG | 119 |
| CDH2 (N-cad) | TGCGGTACAGTGTAACTGGG | GAAACCGGGCTATCTGCTCG | 123 |
| GAPDH | ACAACCTGGTCCTCAGTGTAGCCCA | CATCCACTGGTGCTGCCAAGGCTGT | 220 |

**Supplementary Table S3. DV axis clusters_marker gene list**

**Supplementary Table S4. Neuronal cell type clusters_marker gene list**

**Supplementary Table S5. GEE analysis for morphogenesis by 6AEDs**

**Supplementary video legends**

**Supplementary Video 1. (relate to Fig. 1a). Live imaging of 3D conversion**

After induction of caudal neural stem cells, cell colonies were detached from the dish. Within 12 hrs, colonies were converted to 3D spheroids.

**Supplementary Video 2. (relate to Fig. 1b). Live imaging of the hSCOs exhibiting neural tube closure**

Neural folding-stage hSCOs turned to round spheroid morphology as they completed the neural tube formation-like morphogenesis (Stage II → III).

**Supplementary Video 3. (relate to Fig. 1c). Time-course images of neurulation-like morphogenesis in hSCOs**

Each stage of hSCOs was labeled with ZO1 (white) to visualize the progression of tube-like structure formation.

**Supplementary Video 4. (relate to Fig. 1g). Morphology of hinge cells**

Individual cell morphology was visualized via mixture of GFP-labeled H9 cells. Apical side was labeled with ZO1 (white), and 3D images were processed with Amira software.

**Supplementary Video 5. (relate to Fig. 1l). 3D architecture of neural tube-stage hSCO.**

hSCOs were labeled with SOX2 (green), p-H3 (red), and ZO1 (white). Nuclei were counterstained with Hoechst (blue).

**Supplementary Video 6. (relate to Fig. 3a). Core-to-shell organization of hSCOs**

3-month hSCOs were labeled with NF-M (green), NEUN (magenta) and GFAP (cyan).

**Supplementary Video 7. (relate to Fig. 6e). 3D neural tube morphology in 6 AEDs-treated hSCOs**

3D neural tube morphology was labeled with ZO1 staining after treatments with six different AEDs.
